# Single-cell RNA Sequencing of Pediatric Ependymoma Unravels Subclonal Heterogeneity Associated with Patient Survival

**DOI:** 10.1101/2022.02.26.482082

**Authors:** Haoda Wu, Ruiqing Fu, Yu-Hong Zhang, Zhiming Liu, Zhenhua Chen, Jingkai Xu, Yongji Tian, Wenfei Jin, Samuel Zheng Hao Wong, Qing-Feng Wu

## Abstract

Ependymoma (EPN) is a malignant glial tumor occurring throughout central nervous system which commonly presents in children. Although recent studies have characterized EPN samples at both the bulk and single-cell level, intra-tumoral heterogeneity across subclones remains a confounding factor which impedes understanding of EPN biology. In this study, we generated a high-resolution single-cell dataset of pediatric ependymoma with a particular focus on the comparison of subclone differences within tumors, and show upregulation of cilium-associated genes in more highly differentiated subclone populations. As a proxy to traditional pseudotime analysis, we applied a novel trajectory scoring method to reveal cellular compositions associated with poor survival outcomes across primary and relapsed patients. Furthermore, we identified putative cell-cell communication features between relapsed and primary samples and show upregulation of pathways associated with immune cell crosstalk. Our results reveal both inter- and intratumoral gene expression profiles and tumor differentiation and provide a framework for studying transcriptomic signatures of individual subclones in ependymoma at single-cell resolution.

## Introduction

Ependymomas (EPNs) are primary tumors of the central nervous system that commonly present in childhood. Although the diagnosis and stratification of EPN patients have been facilitated by identification of nine EPN molecular groups from genome-wide DNA methylation studies ^1,2^, EPN patients display a high prevalence of relapse and recurrence typically results in much poorer outcomes ^3^. Between molecular groups, posterior fossa group A (PFA) EPN and supratentorial (ST) EPN with C11orf95-RELA-fusions have been reported to show worse prognoses than PF group B, ST-EPN with YAP1-fusions and spinal EPNs ^1,4^. With the advent of high-throughput single-cell RNA sequencing technologies, recent studies have provided resources for understanding the molecular landscape of EPN, revealing a cellular hierarchy in these tumor cells characterized by an undifferentiated progenitor population which transitions into distinct cell lineages including neuronal precursor-like, glial progenitor-like and ependymal-like cells ^5,6^. However, these studies focused on characterizing EPN from different major molecular groups and anatomical locations, but did not reveal the differences in molecular signatures between subclones within individual tumors.

To date, cellular heterogeneity has typically been viewed as a consequence of hyperproliferation and genomic instability that can give rise to intra-tumoral subclones during tumor progression ^7^. In the context of EPN, previous studies have revealed potential signaling pathways involved in driving the expansion of therapy-resistant EPN subclones, which contribute to tumor relapse and disease progression ^8^. These findings suggest that unravelling subclone-to-subclone variability at single-cell resolution could shed light on the molecular mechanisms underpinning EPN pathogenesis.

In particular, mutant genotypes can grant selective advantage on specific cellular subclones, leading to their outgrowth and allowing them to establish dominance in different types of tissue environments ^7^. To date, cellular heterogeneity has been viewed as a consequence of hyperproliferation and genomic instability that can give rise to intra-tumoral subclones during tumor progression ^7^. For example, genomic instability associated with intra-tumoral heterogeneity can manifest in the form of extensive subclonal evolution demonstrated to be correlated with higher risk of recurrence or death in non-small-cell lung cancer ^9^. Indeed, the emergence of subclonal diversity is a fundamental characteristic of intra-tumoral heterogeneity and has been found to be significantly associated with patient survival across diverse cancer types in a pan-cancer study, including lower-grade glioma and glioblastoma multiforme ^10^. Interestingly, subclones of glioblastoma were shown to display remarkable heterogeneity of drug resistance wherein characteristics of coexisting subclones could be linked to distinct drug sensitivity profiles, hinting at the therapeutic potential of targeted treatments for tumor subclones associated with differential survival outcomes ^11^.

To interrogate subclonal heterogeneity within tumor populations in pediatric EPN, we performed single-cell RNA-seq on EPN samples across PF-A and ST regions. Using a deconvolution approach provided by inferCNV to compare subclones in a single PF-A sample as a proof-of-concept, we identified a subclone-specific cilia-associated program within an individual PF-A EPN sample. We further incorporated a trajectory score analysis to predict correlations between survival outcomes in EPN and molecular characteristics, as well as primary and recurrent tumor populations. Finally, we identified cell-cell communication features between relapsed and primary EPN samples and show upregulation of pathways associated with immune cell crosstalk. Our results reveal gene expression profiles associated with subclonal variability, providing a framework for studies on transcriptomic signatures of brain tumor subclones.

## Results

### Single-cell transcriptomic profiling reveals stemness signature differences between intratumoral subclones in PF-EPN

To characterize intratumoral heterogeneity in human EPN, we performed single-cell RNA sequencing on four EPN patients with the 10x Genomics platform and profiled the transcriptome of 35,102 qualified cells with an average of 3,472 genes per cell (Supplementary Table 1). The cells with high percentage (>12%) of mitochondrial genes and low number (<1500) of genes, and those regarded as doublets were removed from the dataset for subsequent analysis (Supplementary Fig. 1). We first performed copy number variation (CNV) analysis to distinguish neoplastic cells from non-malignant (NM) cells, and identified putative subpopulations of malignant tumor cells with high CNV in each tumor samples (Fig. 1a). Moreover, we applied the approach to score the differentiation state of malignant tumor cells using a panel of differentiation-associated genes, which revealed that CNV-inferred neoplastic cell populations were in a less differentiated state (Fig. 1b).

**Fig. 1.**
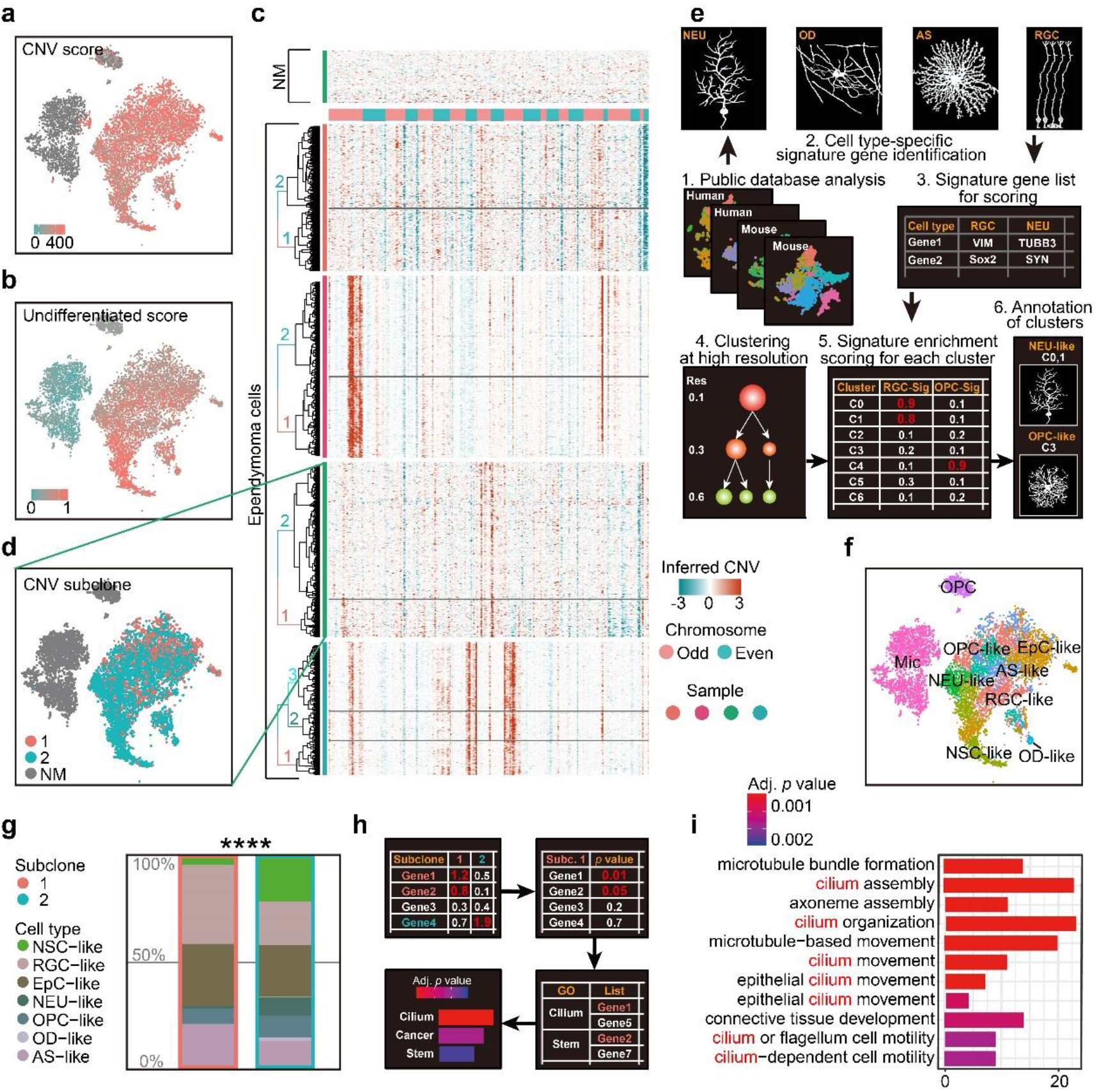
scRNA-seq analysis reveals intratumoral subclone heterogeneity in PF-EPN. **a**, CNV score calculated by modified inferCNV of PF-EPN sample GTE009 presented on tSNE reduction. **b**, Undifferentiated score calculated by CytoTRACE of PF-EPN sample GTE009 presented on tSNE reduction. **c**, CNV heatmap (rows represent cells and columns represent CNV score of genes) of malignant tumor cells from four EPN samples labeled by genetic subclone information for each sample. **d**, Subclonal populations in malignant cells and NM cells of PF-EPN sample GTE009 classified by CNV pattern presented on tSNE reduction. **e**, Workflow of cell-type classification. Signature markers genes are obtained from public transcriptome databases of human and rodent cortex and used for cell-type assignment and cluster annotation. **f**, tSNE plot of all clusters in PF-EPN sample GTE009 color coded by cell types. **g**, Histogram of cell types in PF-EPN sample GTE009 colored by cell-types in percentage and outlined by subclone annotation showing significant difference (*p* value = 7.975e-05) in cell type proportions using asymptotic two-sample Fisher-Pitman permutation test. **h**, Workflow of gene ontology enrichment analysis comparison between PF-EPN sample GTE009 subclone 1 and 2. **i**, Gene ontology analysis of upregulated genes in PF-EPN sample GTE009 subclone 1 compared to the subclone 2 ordered by adjusted p-value.

After subsetting malignant cells from our single-cell dataset, we further examined the presence of different subclones within individual EPN samples. Notably, the inference of subclonal CNV events uncovered the presence of two putative subclones within the PFA-EPN sample GTE009, based on the hierarchical clustering of inferCNV matrix (Fig. 1c–d, and Supplementary Fig. 2). Subsequently, the differentially expressed genes (DEGs) of various cell types, included neural stem cell (NSC), neuron (NEU), radial glial cell (RGC), oligodendrocyte precursor cell (OPC), oligodendrocyte (OD), astrocyte (AS), ependymocyte (EpC), endothelial cells (EC), microglia (Mic), and T cells (TC) from human and rodent embryonic and postnatal cortex scRNA-seq data (Supplementary Table 2), were enriched as signatures to classify cell types in our tumor samples ^12–17^ (Fig. 1e; See Methods). We further applied signature enrichment (SE) analysis and reversed-SE (rSE) for the accuracy of cell-type classification (Supplementary Fig. 3 and Supplementary Fig. 4), which was supported by correlation analysis (Supplementary Fig. 5). Similar to previously published EPN single-cell datasets ^5,6^, our analysis identified NSC-, EpC-, NEU-, RGC-, OPC-, OD- and AS-like cells in malignant populations, as well as endogenous NEUs, ECs, Mic and other cells (Fig. 1f and Supplementary Fig. 4). Although both subclones in GTE009 sample encompassed the same malignant cell types, cell-type composition analysis of these subclones interestingly showed a lower proportion of NSC-like cells in subclone 1 (Fig. 1f–g) with a corresponding increase in EpC-like cells. Moreover, Gene Ontology (GO) analysis revealed enrichment of cilium-related terms based on the DEGs in cells from subclone 1 (Fig. 1h–i) consistent with studies highlighting the role of cilium-related genes in disease processes linked to tumorigenesis ^18,19^, suggesting that molecular characteristics correlated with EPN pathogenesis may be associated with intratumoral subclonal heterogeneity.

### Upregulation of cilium-associated genes is associated with CNV amplification in highly differentiated EPN subpopulations

In spite of transcriptomic and CNV differences between intratumoral subclones, RNAvelocity analysis revealed a classic molecular trajectory originating from NSC-like cells to EpC-like cells similar to previously published single-cell EPN datasets ^5,6^ (Fig. 2a–c). Given the marked differences in cell states between populations from separate intratumoral subclones, we further examined the expression level of differentially expressed cilium related genes and their corresponding CNV score (Supplementary Table 3). We found that cilium-related genes possessed both higher expression and genome amplification in cells from the more highly differentiated subclone 1 compared to subclone 2, suggesting a correlation between CNV amplification and genes associated with more differentiated cell states; for example, the expression level of the cilium-related gene *DYNC2H1*, which has been implicated in the formation of hypothalamic hamartoma ^20^, was both upregulated and amplified in chromosome 11 of subclone 1 (Fig. 2d–f). Indeed, other genes associated with cilium-related terms in GO analysis ^21^ were found to be more highly expressed in subclone 1 compared to subclone 2 (Fig. 2g). The marker genes of EpC-like cells in subclone 1 compared to that of subclone 2 (EpC-Sub1) may be indicative of more mature cellular populations, which was consistent with an inverse correlation with the undifferentiated score (r = −0.65) in EpC-like cells (Fig. 2h–i). These findings suggest that CNV amplification of EpC-related genes is correlated with differentiation of malignant cells, manifesting as alterations in cell composition within intratumoral subclones while maintaining cardinal features of EPN tumorigenesis. This may be relevant in the context of diagnosing malignant EPN samples, as previous findings have shown direct evidence for the roles of CNV-amplified genes in preventing differentiation, inhibiting cell death, and promoting tumor growth, which were in turn correlated with poor patient outcomes ^22^.

**Fig. 2.**
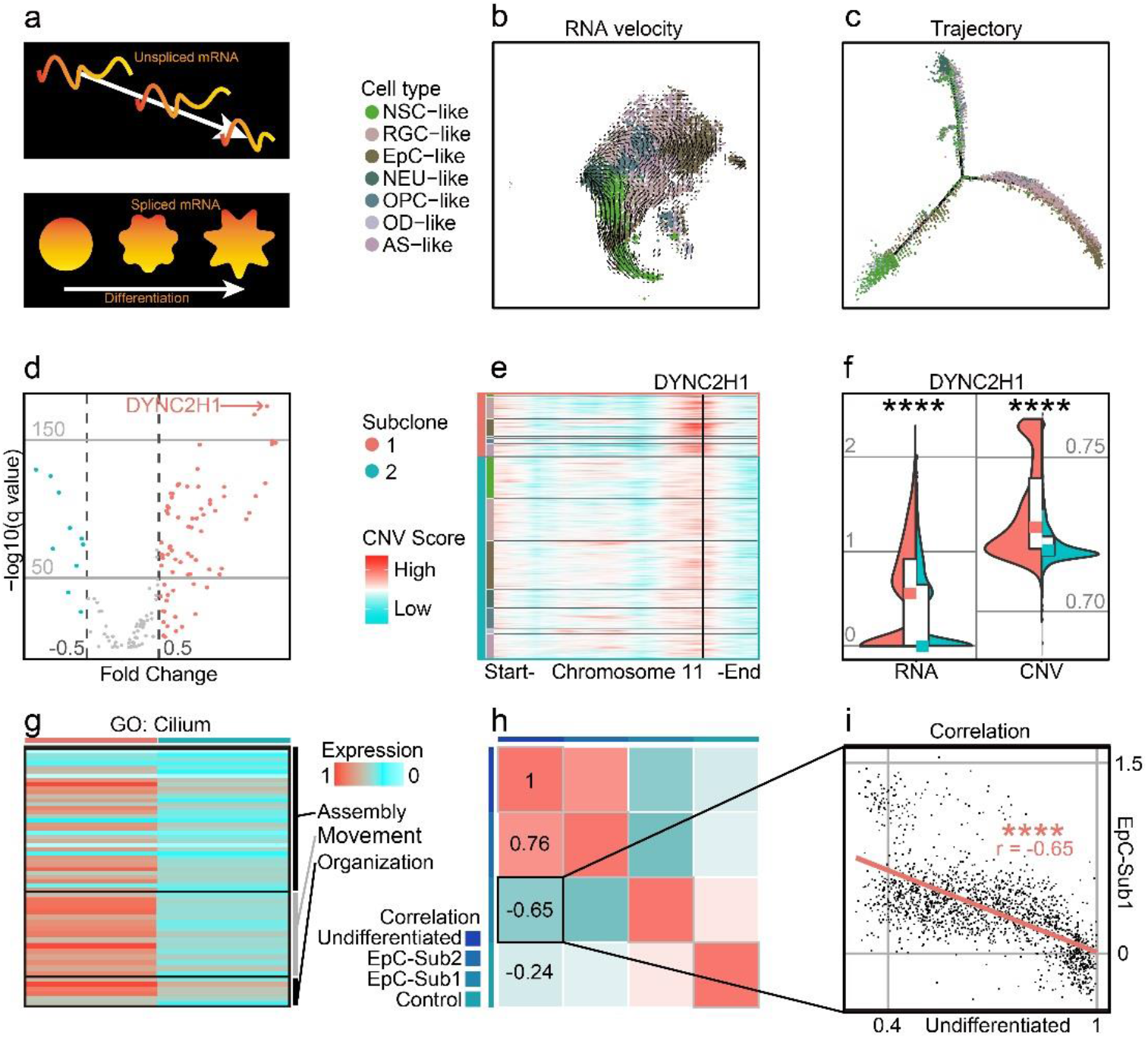
Highly differentiated cells in PF-EPN subclonal populations show CNV amplification and enrichment of cilium-associated genes. **a**, Schematic of RNA splicing analysis and cell differentiation using RNA velocity and trajectory deduction methodologies. **b**, RNA velocity inferred by Velocyto and scVelo of malignant tumor cells presented on tSNE reduction and colored by cell types in PF-EPN sample GTE009. **c**, Differentiation trajectory inferred by Monocle of malignant tumor cells in PF-EPN sample GTE009. **d**, Volcano plot showing genes with differentially expressed CNV values highlighting *DYNC2H1* in PF-EPN sample GTE009 subclone 1 compared to the subclone 2. **e**, Heatmap of chromosome 11 showing inferCNV scores colored by cell types designated in **Fig. 2b** and subclone annotation in PF-EPN sample GTE009. *DYNC2H1* is highlighted by black vertical bar. **f**, Violin plot showing significant difference (*p* value < 0.0001; Mann Whitney test) in gene expression (RNA) and CNV level of *DYNC2H1* between subclones in PF-EPN sample GTE009. **g**, Heatmap showing relative expression of identified genes from cilium-related terms in GO analysis ^21^ colored by subclones in PF-EPN sample GTE009. **h**, Correlation analysis of undifferentiated score in EpC-like cells, normalized average expression of markers in EpC-like cells in subclone 1 (EpC-Sub1) and subclone 2 (EpC-Sub2), and normalized average expression of Mic (Control; see Supplementary Table 2) in PF-EPN sample GTE009. **i**, Pearson correlation between undifferentiated score and normalized average expression of EpC-Sub1 (*p* value < 0.0001) in PF-EPN sample GTE009.

### Trajectory score analysis identifies cellular compositions associated with worse survival outcomes in EPN

To further analyze the cellular subpopulations in subclone 1 and 2, we performed GO analysis on the DEGs between subclones 1 and 2 in NSC-like and EpC-like cells, respectively. We detected high expression of cell-cycle-related genes in NSC-like cells from subclone 2, as well as enrichment of cilium-related genes in EpC-like cells in subclone 1 (Fig. 3a). Given that previous studies have demonstrated that high expression of NSC-like cells and low expression of EpC-like cells are correlated with poor patient survival, and vice versa ^5,6^, we hypothesized that performing a combinatorial analysis integrating information from both subclone and cell-types may help in predicting EPN patient outcomes.

**Fig. 3.**
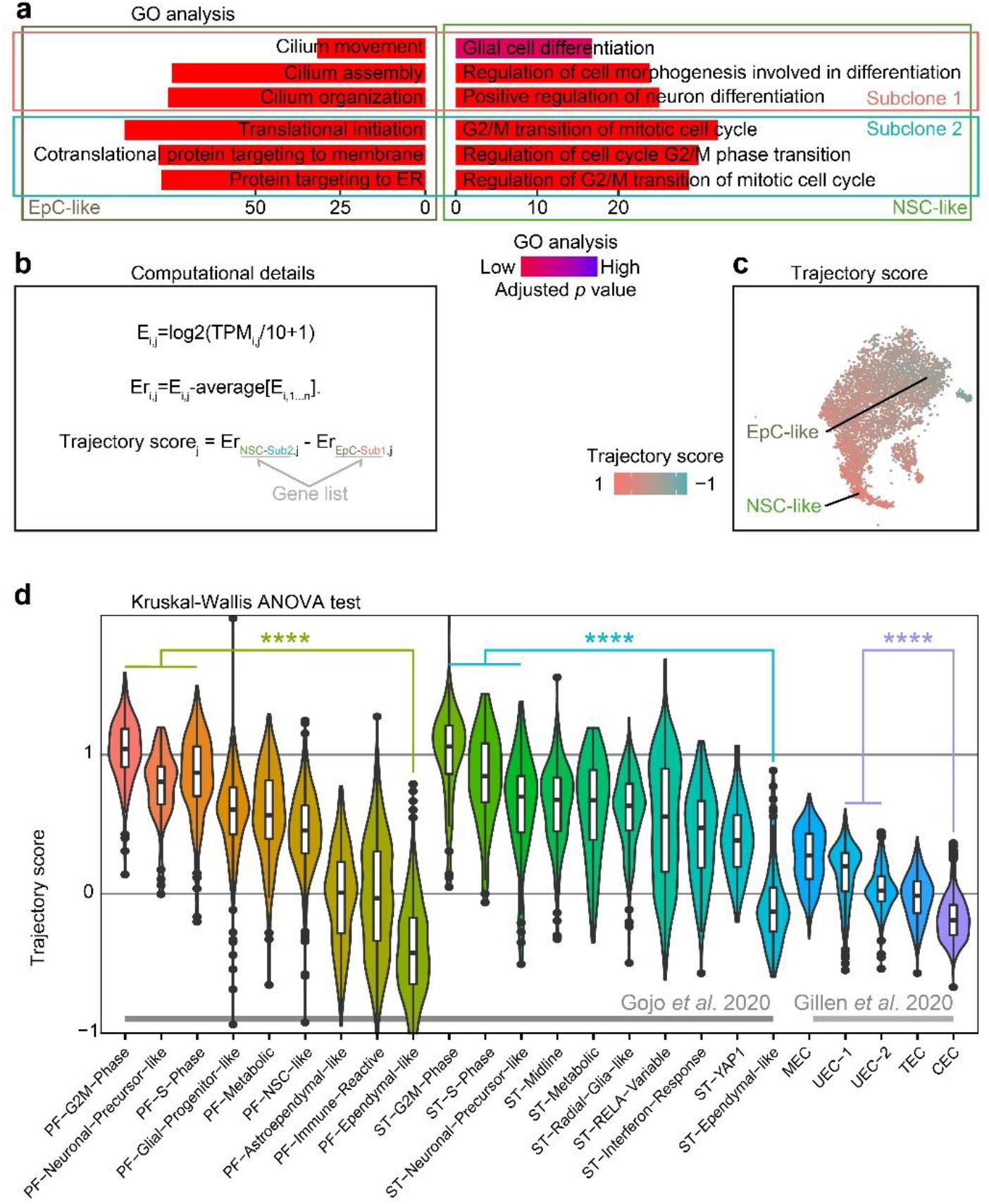
Trajectory score analysis can predict EPN cell compositions associated with poor survival outcomes. **a**, Gene ontology analysis of differentially expressed genes from EpC-like and NSC-like cells between subclones classified by annotation of subclones and cell types in PF-EPN sample GTE009. **b**, Workflow for calculating trajectory score based on published computation method ^35^. E: expression; TPMi,j: transcript-per-million (TPM) for gene i in sample j; Er: relative expression. **c**, tSNE plot of trajectory score in combined subclone 1 and 2 datasets with EpC-like and NSC-like cell populations labeled in PF-EPN sample GTE009. **d**, Validation of trajectory score on published scRNA-seq data ^5,6^ of EPN using previously defined cell-type annotations (*p* value < 0.0001; Kruskal-Wallis test).

To provide a numerical representation of molecular trajectory information at the single-cell level, we developed a trajectory score for downstream analyses (Fig. 3b): the normalized average expression of undifferentiated-NSC (NSC-like cells in the subclone 2; NSC-Sub2) markers was subtracted by differentiated-EpC (EpC-Sub1) markers. The trajectory score was presented in tSNE plot (Fig. 3c), which resembled the trajectory analysis (see Fig. 2b). The trajectory score allows for identification of differentiation state at the single-cell level which complements molecular trajectory analysis in merged samples. Notably, well-characterized stemness-associated markers ^10,23^ such as *FTL*, *LGALS1*, *MEG3*, *MEST*, *TUBB*, *TMSB4X*, and *STMN1* were found in the DEGs of trajectory-high group compared to trajectory-low group (Supplementary Table 4), while cilium-related terms were enriched in the trajectory-low group compared to trajectory-high group (Supplementary Fig. 6a).

Based on the hypothesis that trajectory score was directly correlated with survival outcome and could be easily used to predict the prognosis of EPNs by its simple calculation method, we applied this scoring methodology to two published EPN scRNA-seq datasets (33 EPN patients in total), and show that our results are consistent with the respective survival outcomes reported in these studies based the respective cell-type composition of individual samples ^5,6^ (Fig. 3d). On the contrary, application of the aforementioned undifferentiated score led to relatively more inconsistent results (Supplementary Fig. 6b), suggesting that trajectory score analysis could be a useful tool to investigate EPN prognosis. Indeed, samples with a high trajectory score were found to have correspondingly poorer survival outcomes in published PF-EPN samples and PF/ST-EPN ^5,6^ (Supplementary Fig. 6c), although the comparison did not reach significant difference due to the small sample size. Likewise, a higher percentage of recurrent patients compared to primary patients had higher trajectory score (Supplementary Fig. 6d), supporting the association between EPN relapse state and overall survival outcomes of this disease.

### Cell compositions correlated with poor prognosis in EPN recurrent patients are revealed by trajectory score analysis

Relapse rates for EPN can be as high as one third of patients ^3^, and relapse is known to lead to significantly worse survival outcomes based on published data ^5,6^ (Fig. 4a). To determine transcriptomic signatures at the single-cell level associated with poor survival outcomes in EPN relapse, we compared the cell-type composition in 36 EPN patients ^5,6^ and revealed a significant difference with higher percentage of NSC-like cells in relapse patients (Fig. 4b). Similarly, we found a significantly higher trajectory score in recurrent NSC-like cells compared to primary NSC-like cells, and a similar trend was observed for EpC-like cells (Fig. 4c). Given the association between high stemness and poor survival outcome found in relapse patients based on the trajectory score, we further performed GO analysis on the DEGs between NSC-like cells in primary and recurrent patients and revealed enrichment of cilium- and immune-related terms (Fig. 4d). This suggests that NSC-like cells from recurrent patients were not only in a more immature cell state, but also that there is a likelihood of extensive immune cell crosstalk within these cellular populations, consistent with previous findings reporting the association between cell-cell communication in immune cells and tumor progression ^24,25^.

**Fig. 4.**
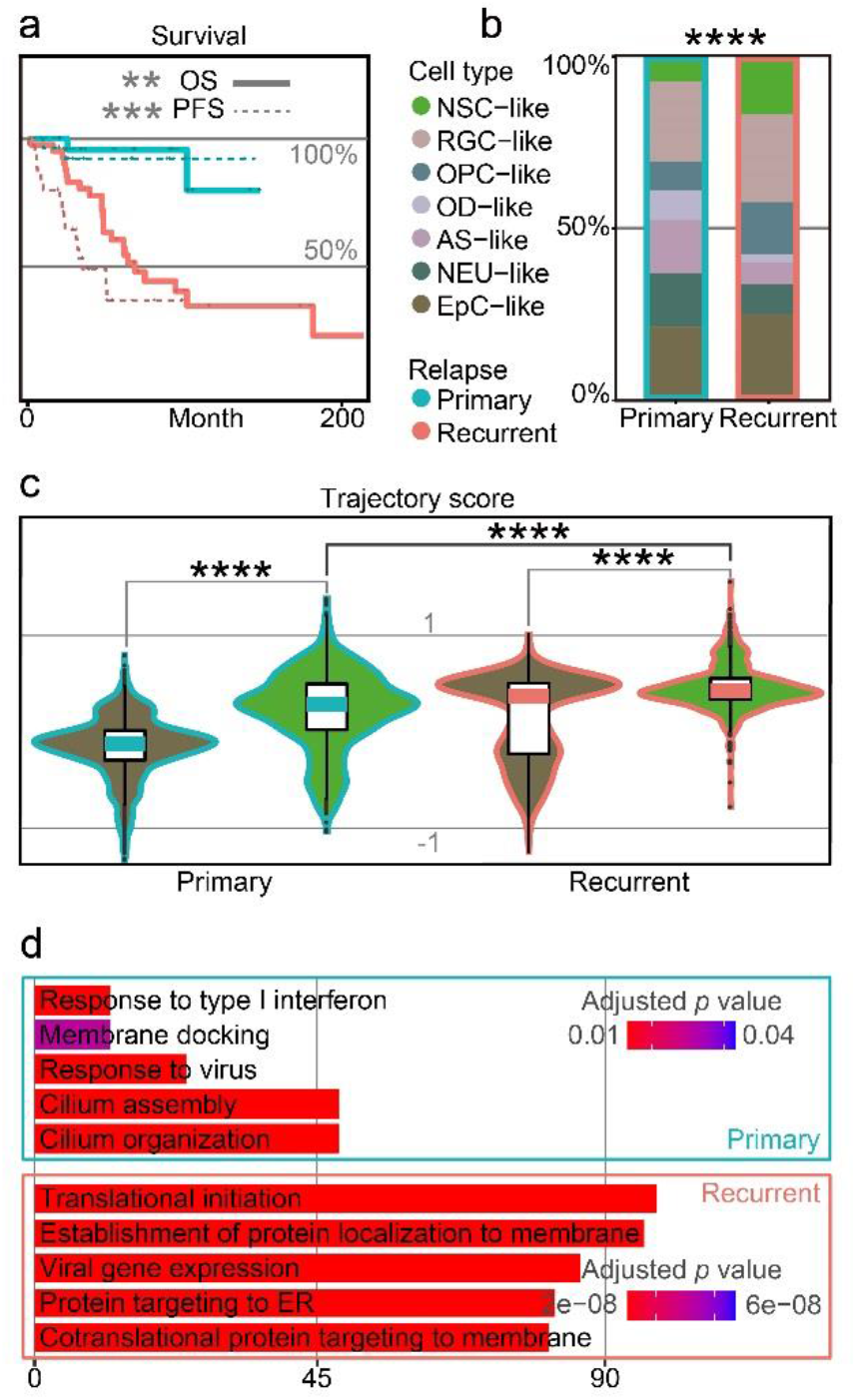
Cellular populations in recurrent EPN with poor prognosis are associated with higher trajectory score. **a**, Survival plot of primary and recurrent EPN patients. The solid line refers to overall survival (OS; *p* value = 0.0018) and the dotted line refers to progression-free survival (PFS; *p* value = 0.00026), which are colored by relapse situations on published scRNA-seq data ^5,6^. **b**, Histogram of cell types in primary and recurrent EPN colored by cell-types and outlined by primary/recurrent conditions showing significant difference (*p* value < 2.2e-16) between cell types using asymptotic two-sample Fisher-Pitman permutation test. **c**, Trajectory score analysis comparison between primary and recurrent samples in NSC-like and EpC-like cells using Kruskal Wallis test (all *p* values < 0.0001). **d**, Gene ontology analysis of differentially expressed genes in NSC-like cells between primary and relapse conditions.

### Relapsed EPN show upregulation of distinct signaling pathways associated with immune cell crosstalk

Based on studies demonstrating the role of tumor-infiltrating NM cells such as Mic in brain tumor ^24,25^, we performed crosstalk analysis to investigate cell-cell interactions between the different cell-types profiled. Although there has been increasing interest in examining communication patterns between cell populations using scRNA-seq, crosstalk analysis on intracranial EPN samples has not been extensively studied given the relatively lower number of tumor-infiltrating NM cells profiled in previous datasets ^5,6^. To investigate cell– cell interactions in EPN, we first examined the expression of ligand–receptor pairs in different cell-types across four EPN samples (Fig. 5a) and identified the presence of numerous crosstalk events (Fig. 5b, Supplementary Fig. 7a and Supplementary Table 5). For example, we observed strong outcoming events from NSC-like cells towards other cells (Supplementary Fig. 7b). Moreover, we uncovered the overlap in simulated spatial 3D position between NSC-like cells and Mic (Fig. 5c and Supplementary Fig. 7c). Interestingly, these events that had higher expression in recurrent samples than that in primary samples in 36 EPN patients (Fig. 5d) shown interaction between NSC-like cells and Mic (see Supplementary Fig. 7d–e), including MK pathway that promoted brain tumor growth ^24,25^ and *EGFR* pathway that inhibited glioblastoma invasion via pharmacological inhibition of *EGFR* ^26^. For example, *MDK* (ligand) and *NCL* (receptor) were highly expressed in the tumor microenvironment (TME) of recurrent samples (Fig. 5e), consistent with previous studies implicating the roles of this ligand-receptor pair in tumorigenesis ^27,28^ and the MK-deficiency reduced tissue infiltration of microglia ^29^. To further elucidate signaling pathways involved in crosstalk between normal and malignant cells, the inferred gene regulatory networks also revealed multiple pathways shared in crosstalk (see Supplementary Results and Supplementary Fig. 8). Taken together, crosstalk analysis on 35,102 individual cells in conjunction with validation using 36 EPN patients revealed elevated cell-cell interactions between malignant cells and tumor-infiltrating NM cells, such as between NSC-like cells and Mic, consistent with studies demonstrating a key role of the central nervous system TME in the pathogenesis of EPN.

**Fig. 5.**
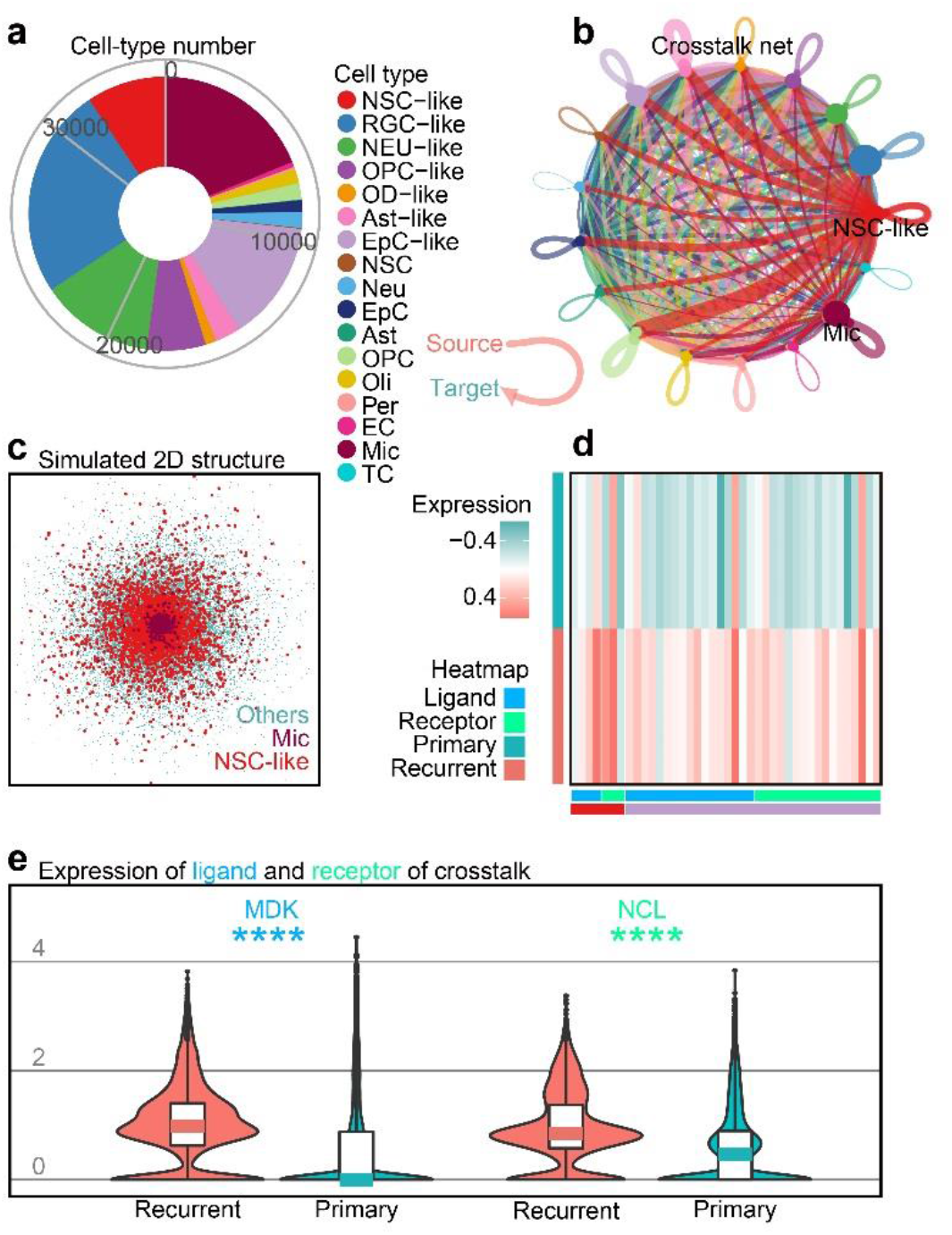
Crosstalk analysis reveals cell-cell interactions implicating immune cell populations in recurrent EPN. **a**, Cell numbers of all cell-types in four EPN samples. **b**, Crosstalk net analyzed by CellChat. Individual lines represent the crosstalk from source to target cells. Related to Supplementary Fig. 7a–b. **c**, Simulated 2D spatial structure showing overlap of Mic and NSC-like cell populations by CSOMAP. Related to Supplementary Fig. 7c. **d**, Heatmap of ligands or receptors with significantly higher expression in recurrent samples compared to primary samples, colored by cell type and gene class (ligands or receptors) using published single-cell transcriptomes of 36 EPN samples ^5,6^. **e**, Expression of *MDK* (ligand) and *NCL* (receptor) between recurrent and primary samples of 36 EPN patients ^5,6^ (*p* value < 0.0001; Mann-Whitney test).

## Discussion

The increasing accessibility of scRNA-seq technologies has accelerated our understanding of cellular function in health and disease. Here, we generated a high-resolution EPN single-cell dataset with a particular focus on the comparison of subclone differences within tumor populations. Our analysis on four EPN samples profiled 35,102 single-cell transcriptomes and uncovered 17 major cell types including NSC-like, EpC-like and microglia populations that are present across different EPN groups. We further reveal differences in cell proportions within highly differentiated populations within tumor subclone cells by integrating CNV pattern analysis with single-transcriptome data in this study and previously published datasets. Additionally, differential gene expression analysis also identified gene programs associated with tumor subclones as well as survival outcomes.

Treatment of heterogeneous tumors such as EPN can favor selection of resistant subclones given that different subclones respond differently to intrinsic and extrinsic signaling cues. EPN relapses after surgical resection and treatment of EPN are common and have poor outcomes, and recent findings have demonstrated that administration of radiation and chemotherapy can lead to a significant increase in EPN mutational burden in conjunction with changes to the tumor subclonal architecture, without eliminating the original founding clone ^8^. In this study, we report the presence of subclones within a single EPN tumor sample characterized by molecular signatures reflecting different stages of cellular differentiation. This suggests that in addition to stemness signature gradients between tumors ^5,6^, intratumoral heterogeneity can also be uncovered within individual samples containing multiple subclones. Importantly, further interrogation of the subclones identified from CNV analysis also showed that EPN subpopulations which were more differentiated exhibited an increase in cilium-associated genes. For example, in the sample GTE009, 6% of 494 DEGs from EpC-like cells compared to the other malignant tumor cells were found to be overlapped with genes related to cilium assembly, organization, and movement such as *PIFO, ZMYND10, DNAH9, TEKT1, DYNLL1, SPEF1, MNS1, DNAAF1, IQCG, SPAG16, FOXJ1*. Indeed, ciliary signaling is known to be a mediator of paracellular signals controlling cancer metastatic processes and responses to therapy, and mutations leading to defects or structural abnormalities in cilia have been shown to be directly correlated with cancer pathogenesis^18^. Given that ependymal cells in EPN are multiciliated cells ^19^, it is plausible that changes in ciliation of EPN subpopulations and/or cells of the tumor TME during EPN development can contribute to disparities in outcomes within tumors of the same molecular group.

To complement classical pseudotime molecular trajectory methodologies ^71^, we applied a curated trajectory score to our single-cell dataset to integrate EPN subclone and cell-type information in our analysis and found that higher trajectory scores were found in patients with EPN samples exhibiting elevated stemness signatures (*FTL*, *LGALS1*, *MEG3*, *MEST*, *TUBB*, *TMSB4X*, *STMN1* ^10,23^), which is also associated with worse prognoses. This trajectory score analysis allows for a numerical representation of EPN trajectory stages at the single-cell level to facilitate comparison with datasets of interest. Moreover, our analysis takes into consideration both cell-types and stemness signatures and can be easily applied to other transcriptomic datasets of interest to quantify the relative correlation degree between survival outcomes and stemness signatures across different samples. As a proof-of-concept, we performed validation of our dataset with previously published results on EPN samples (derived from 36 patients in total), and indeed demonstrated that this analysis reveals consistent trends in EPN survival outcomes. In addition to using trajectory score analysis to examine the association between EPN stemness signatures and survival outcomes, we further applied this method to compare differences in survival outcomes between primary and recurrent EPN samples. Previous findings have identified an enrichment of undifferentiated programs (NSC-like) in recurrent PF-EPN relative to primary PF-EPN samples from comparing three matched samples at the single-cell level ^5^. Here, we find that this is consistent across the single-cell transcriptome of 36 published EPN samples, as trajectory score analysis shows a clear significant difference in cell-type composition between recurrent and primary EPN samples with a higher percentage of NSC-like cells in recurrent EPN. Further analysis of subpopulations within recurrent and primary samples revealed that recurrent samples also show higher trajectory score in NSC- like cells and in EpC-like cells compare to the corresponding cell types in primary samples. These findings suggest that trajectory score analysis can uncover multiple types of association in EPN samples in a quantifiable form, such as the correlation between stemness and tumor occurrence with patient mortality.

In EPN and other brain cancers, there is increasing evidence that the brain TME functions as a key regulator of cancer progression in brain malignancies ^30^. Hence, we performed cell-cell communication analysis on our EPN single-cell transcriptomes to assess the crosstalk between different cell types in EPN, given that the TME contains non-cancerous cell types such as pericytes, endothelial cells, and immune cells in addition to cancer cells. We revealed putative interactions between malignant cells and tumor-infiltrating NM cells, such as enrichment in interactions between NSC-like cells and microglia. For example, the MK pathway, which has been implicated in brain tumor pathogenesis ^24,25^, was not only found to be a significantly upregulated in this study’s dataset, but also showed higher expression in recurrent samples compared to primary samples based on 36 published EPN single-cell transcriptomes. Indeed, MK deficiency has been shown to reduce tissue infiltration of microglia, leading to reduced neuroinflammation and apoptosis ^29^. Given that inflammatory cross-talk with immune cells has previously been shown to play a key role in driving tumor growth in the EPN microenvironment ^31^. These findings suggest that immune cell crosstalk analysis may serve as a useful resource for identification of candidate genes for future in vitro and in vivo validation studies.

In summary, we report a curated EPN atlas focusing on comparison of intra-tumoral heterogeneity in this study. We use an integrative analysis approach to show both changes in cell-type composition and cell-type-specific gene expression associated with different tumor groups and subclones. Moreover, we also apply a novel trajectory scoring method as a parallel tool to traditional molecular trajectory analysis and demonstrate its robustness in recapitulating survival outcomes within individual EPN samples and across primary and recurrent tumors. This approach will complement existing published datasets and provide valuable insights into cell-type-specific properties of EPN, laying the foundation for therapeutic treatments of this disease.

## Materials and Methods

### EPN sample preparation for scRNA-Seq

Fresh tumor samples were processed as previously described with minor modifications ^32^. Fresh EPN tissue was excised by physicians with signed informed consent documents that was approved by the ethics committee of Beijing Tiantan Hospital of Capital Medical University. Samples were delivered on ice to Institute of Genetics and Developmental Biology of Chinese Academy of Sciences immediately. Microdissected tissues were transferred to a 24-well cell culture plate and digested by buffer comprising 20 U/mL Papain (LK003178; Worthington, Lakewood, U.S.A.), 100 U/mL DNaseI (LK003172; Worthington), 10 U/L chondroitinase ABC (C3667; Sigma-Aldrich, St. Louis, U.S.A.), 0.07% hyaluronidase (R006687; Rhawn, Canton, P.R.C.), 1 X Glutamax (35050061; Life Technologies, Waltham, U.S.A.), 0.05 mM (2R)-amino-5-phosphonovaleric acid (APV; 010510; Thermo Fisher Scientific, Waltham, U.S.A.), 0.01 mM Y27632 dihydrochloride (T9531; Sigma), and 0.2 X B27 supplement (17504044; Thermo Fisher Biosciences) in Hibernate-E media (A1247601, Life Technologies) for 1-2 hrs at 37°C, and then pooled with Hibernate-E buffer containing 1xGlutamax, 0.05 mM APV, 0.2 X B27, 0.01 mM Y27632 dihydrochloride. Tissues were gently triturated through Pasteur pipettes with finely-polished tips of 600, 300 and 200 μm diameters in order, and washed once with Hibernate-E buffer to generate single-cell suspensions. After filtration through a 40 μm strainer (130-101-812; Thermo Fisher Scientific), 1 X red blood cell lysis solution (130-094-183; Miltenyi Biotec, Bergisch Gladbach, Germany) was added to remove blood contamination during surgery followed by 1800 μL debris removal solution (130-109-398; Miltenyi Biotec). Subsequently, the dissociated cells were stained with DAPI (0.2 μg/mL) to identify dead cells. scRNA-seq libraries were constructed under the manufacturer instructions provided by 10x Genomics accompanying single cell 3′ Library and Gel Bead Kit V3 (1000075; 10x Genomics, Pleasanton, U.S.A.). A Chromium Single Cell Controller (10x Genomics) loaded cell suspensions (300-600 living cells per microliter determined by Count Star) to generate single-cell gel beads in the emulsion (GEM). Quality control was performed on the generated cDNA library using the Agilent 4200 and scRNA-seq was performed on the Illumina Novaseq6000 sequencer.

### Data processing of scRNA-seq

The sequencing data was processed by CellRanger v3.1.0 with reference genome hg19-3.0.0 to generate filtered expression matrices which were analyzed using Seurat v3.2.0 ^33^. Doublet Finder ^34^ was first applied to erase doublets with default settings. Genes detected in at least ten cells were used for analysis, and cells that possessed transcription numbers fewer than 1,500 or cells with mitochondrial genes talking up more than 12% of reads were removed. After normalizing the data, we used 5,000 highly variable features for downstream analysis and cell cycle variation was regressed out as previously described ^35^. tSNE analysis was performed with top 50 significant principal components from principal component analysis and cells were clustered using ‘FindClusters’ function based on tSNE reduction. CNV analysis was performed using inferCNV of the Trinity CTAT Project^36^ (https://github.com/broadinstitute/inferCNV) as described in the following section. Calculation of the undifferentiated score was performed using CytoTRACE ^37^ in as previously described ^32^. The EPN single-cell datasets covering malignant cells and tumor-infiltrating NM cells were analyzed by ‘iCytoTRACE’ function as the quantification of the number of expressed genes as an indicator of differentiation potential; genes associated differentiation (deduced by CytoTRACE) and their relative expression levels were used to perform this calculation. KEGG/GO analysis was performed based on the DEGs between malignant tumor cells and NM cells of the merged dataset using ‘FindAllMarkers’ function in Seurat and clusterProfiler ^38^. The correlation of malignant tumor cells and NM cells among samples was in favor of malignancy separation (Supplementary Fig. 1e) utilizing ‘cor’ function of stats package in R v3.6.3.

### Estimating CNVs in scRNA-seq data

The initial CNVs of single cells were estimated from their whole-genome wide expression level by inferCNV ^35,36^. To perform comparison across samples, 300 cells of OPCs were sampled from the GTE009 as a common reference, and all the non-immune cells of each sample were tested against it, with the parameters of ‘min_max_counts_per_cell = c(5e2, 6e6); cutoff = 0.1; min_cells_per_gene = 5’ and other default parameters in inferCNV. Then, the estimated CNV values were re-scaled to 0 to 2, with 1 as the normal copy, to compare among samples. The CNV cluster of each sample were deduced by the hierarchical clustering (ward.D2) of inferCNV matrix. The CNV level of each cell was also calculated as previously described ^39^. The estimated CNV values were re-standardized as −1 to 1, and the CNV level of each cell was then calculated as the quadratic sum of all the expressed genes.

### Estimating CNVs in WES data

Sample DNA was extracted and sequenced via Agilent SureSelect Human All Exon v6 and Illumina platform. Whole-exome sequencing reads were aligned to human reference genome (b37), using BWA ^40^, followed by marking of duplications via Picard (http://broadinstitute.github.io/picard/). CNVkit ^41^ was used to call CNVs from targeted regions of exons in each sample, following the default workflow, with the bin size of 1kb. A flat reference was made to run CNVkit with each sample, and during segmentation, the ‘CBS’ method was applied, with 1e-4 as the significance threshold and parameters of ‘-- drop-low-coverage --drop-outliers 3’.

### Statistical analysis

CNV score, gene expression, and undifferentiated score comparison between subclones was analyzed by D’Agostino & Pearson normality test, Shapiro-Wilk test, and Mann Whitney test in GraphPad v8.3.0. Asymptotic Two-Sample Fisher-Pitman Permutation Test of cell-type composition between subclones of the sample GTE009 and between recurrent and primary samples was performed by ‘oneway_test’ function of coin package in R v3.6.3. Correlation analysis was performed by R package ‘corrplot’ from Taiyun Wei and Viliam Simko (2021; : Visualization of a Correlation Matrix (Version 0.90)) and cor() function in R v3.6.3.

The survival analysis was performed on published data ^5,6^ by R package ‘survminer’ from Alboukadel Kassambara, Marcin Kosinski and Przemyslaw Biecek (2021; Drawing Survival Curves using ‘ggplot2′ (0.4.9)) and R package ‘survival’ ^42^. The groups were separated by the relapse situation (recurrent or primary annotated by original authors ^5,6^) or the mean of trajectory scores of samples. First, a survival object was created by ‘Surv’ function; second, the survival curves were created by ‘survfit’ function based on a tabulation of the number at risk and at death time of events from the supplementary files of these published researches ^5,6^; third, ‘ggsurvplot’ function was used for the visualization of these curves.

### Cell-type annotation

For cell malignancy analysis, we combined the following approaches to achieve a combinational separation of non-malignant cells and malignant tumor cells. First, all cells were sorted by t-distributed stochastic neighbor embedding (t-SNE) projections in each patient and colored based on the cell clusters identified by Seurat ^33^, as malignant cells were often comprised of multiple clusters and were contiguous in t-SNE projection ^43^. Second, the malignancy of cells was explored by copy number variation (CNV) scores through whole-exon sequence by CNVkit ^41^ (Supplementary Fig. 2) and through modified inferCNV (https://github.com/broadinstitute/inferCNV) ^35,36^ (Fig. 1a) for each sample. Third, the malignancy was further supported by the high undifferentiated score calculated by CytoTRACE ^37^ (Fig. 1b). Combining the result of cell-cycle stages (Supplementary Fig. 1d), we classified each sample into malignant tumor cells and non-malignant cells. For the better exploration of the intratumoral genome, the pattern revealed by inferCNV was used for separation of malignant tumor cells into different subclones of each sample (Fig. 1c–d). To support the genetic information inferred by transcriptome, we applied whole-exome sequencing analyzed by CNVkit ^41^ (Supplementary Fig. 2) and obtained similar results in chromosomal level comparing to that from inferCNV. Fourth, the high correlation between malignant tumor cells and between non-malignant cells among samples supported malignancy separation (Supplementary Fig. 1e). As validation, we performed Kyoto encyclopedia of genes and genomes (KEGG) analysis ^38^ on the differentially expressed genes (DEGs) of malignant tumor cells compared to non-malignant cells of merged samples, which shown enrichment on cell cycles and cancer-related terms: breast cancer, hepatocellular carcinoma, and proteoglycans in cancer (Supplementary Fig. 1f). To sum up, we obtained non-malignant cells and malignant tumor cells and explored the genome of each sample.

To achieve cell type classification, we applied signature enrichment analysis (Supplementary Fig. 3). Firstly, the DEGs of various cell types were calculated from each published data of corresponding area (Supplementary Table 2) and were used as signatures to distinguish cell types in our tumor samples ^12–17^ (Fig. 1e) which included RGC, AS, EC, EpC, NEU, NSC, OD, OPC, Mic, and T cells, from human and rodent embryonic and postnatal cortex scRNA-seq data. The enrichment of these gene signatures was calculated using the ‘AddModuleScore’ function by subtracting the aggregated expression of control genes from the average expression levels of gene signatures ^35^. Then the signature enrichment of those gene sets in cellular level was summarized in cluster level on average and was utilized as the standard to ascertain the cell type. Conclusively, the cell types of clusters were determined by the highest signature score.

Studies on scRNA-seq have also employed a reversed approach for cell type classification (rSE; extracting signatures from unknown clusters and enriching them on published data of identified cell types; for details, see methods; Supplementary Fig. 3) ^5,6,44,45^. We then compared the similar results of these two methods in our data and obtained a high degree of correlation (Supplementary Fig. 3). Hence, the following research applied the result of the first mentioned method, SE analysis. This method and calculated signatures (Supplementary Table 2) were further tested on other published data, which displayed high similarity in cell type classification compared to the cell types determined by original authors: human cortex ^46^ (Supplementary Fig. 5b), rodent cortex ^17^ (Supplementary Fig. 5c), ependymoma ^5^ (Supplementary Fig. 5d), and childhood ependymoma ^6^ (Supplementary Fig. 5e).

### Trajectory analysis

For developmental trajectory analysis, the BAM files from Cell Ranger were processed by Velocyto ^47^ to obtain loom files containing spliced and unspliced transcript counts, which was used as input for scVelo ^48^ and Velocyto ^47^. Monocle ^49,50^ was applied on 2000 variable features detected within more than 5% of cells to obtain reduced coordinates by ‘DDRTree’. The trajectory score was inferred by ‘AddModuleScore’. The undifferentiated trajectory score was calculated based on the significantly highly expressed genes in NSC-like cells in subclone 2 of the sample GTE009, while the differentiated trajectory score was calculated based on the significantly highly expressed genes in EpC-like cells in the subclone 1 of the sample GTE009. The final trajectory score was then calculated by subtracting the differentiated trajectory score from the undifferentiated trajectory score of each cell.

### Crosstalk and gene regulatory network analysis

Crosstalk analysis on EPN was performed on the integrated dataset of four EPN samples using CellChat ^51^ and simulated 3D spatial structure of different cell types was calculated by CSOMAP ^52^. In details, CellChat preprocessed the expression data of our integrated four ependymoma for cell-cell communication analysis; the cell-cell communication network was inferred by computation of the communication probability at a signaling pathway level and the calculation of the aggregated data frame. The elevated crosstalk pathways were validated in previously published EPN single-cell datasets ^5,6^. Similarly, CSOMAP computed the network of ligand-receptor interaction and then calculated optimized 3D coordinates.

Regulons for individual cell types were computed using the SCENIC (single-cell regulatory network inference and clustering) pipeline ^53^ on our integrated four EPN samples and validated by previously published EPN single-cell datasets ^5,6^. A log-normalized expression matrix of the four integrated EPN samples was used as an input into the pySCENIC workflow with default settings to infer regulon activity scores. To examine relevant networks using cell-cell communication analysis, we identified genes involved in crosstalk (ligands or receptors expressed in NSC-like cells) or gene regulatory networks (regulons with significantly high activity in NSC-like cells) of interest and then used genes classified under the same enriched terms in GO/KEGG analysis. Genes that had significantly higher expression in recurrent EPN than that in primary EPN were enriched in GO and KEGG analysis to highlight key terms in crosstalk and regulatory networks.

## Acknowledgements

The work was supported by the National Key R&D Program of China (2019YFA0801900 and 2018YFA0801104), the National Natural Science Foundation of China (81891002, 31921002, 32070972 and 31771131), the Strategic Priority Research Program of Chinese Academy of Sciences (XDB32020000) and the Beijing Municipal Science & Technology Commission (Z210010 and Z181100001518001).

## Ethics declaration

### Competing interests

The authors declare no competing interests.

## Supplementary information

Supplementary Figures 1–8

Supplementary Tables 1-5

## Supplementary information

**Supplementary Fig. 1.**
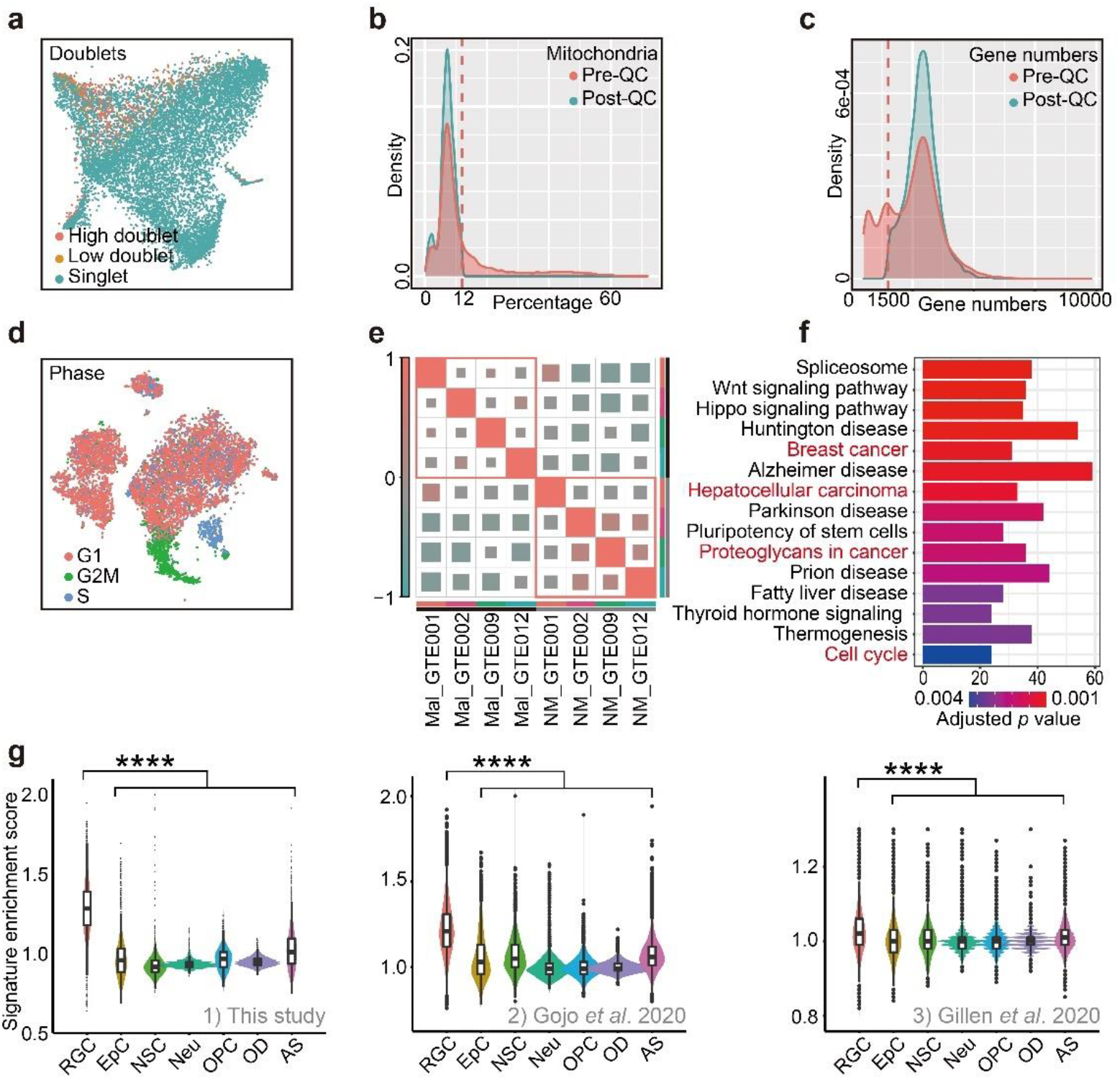
Quality Control of scRNA-Seq Analysis of Human EPN. **a**, Doublets identified by Doublet Finder presented on tSNE reduction. High doublet cells are filtered out. **b**, Density plot of mitochondria percentage of cells in sample GTE009. Cells with more than 5% of mitochondria percentage are filtered out. **c**, Density plot of captured gene numbers of cells in sample GTE009. Cells with less than 1500 transcripts are filtered. **d**, Cell cycle phases of cells in sample GTE009 re-calculated after quality control presented on tSNE reduction. **e**, Correlation plot of transcriptome of malignant tumor cells (Mal) and non-malignant cells (NM) among samples (GTE001, GTE002, GTE009, and GTE012). **f**, KEGG analysis on DEGs of malignant tumor cells compared to non-malignant cells of four merged samples (GTE001, GTE002, GTE009, and GTE012). **g**, Enrichment of RGC signatures in malignant tumor cells compared to other cells types using single-cell transcriptomes from 1) this study, 2) EPN ^5^ and 3) childhood EPN ^6^ (one-way ANOVA analysis; p value < 0.0001).

**Supplementary Fig. 2.**
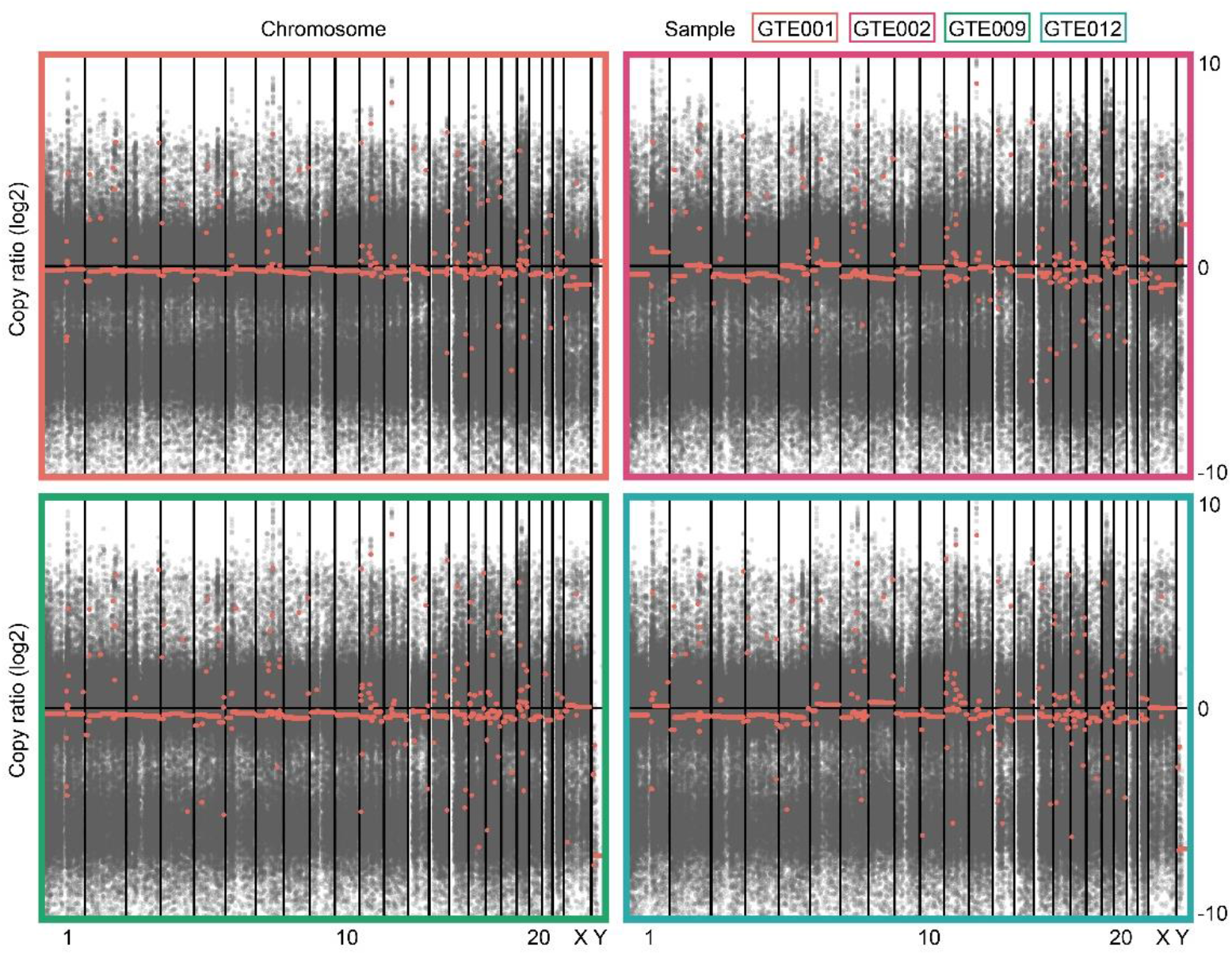
CNV Analysis of Whole-exome Sequencing. CNV heatmap of whole-exon sequence data labeled by samples.

**Supplementary Fig. 3.**
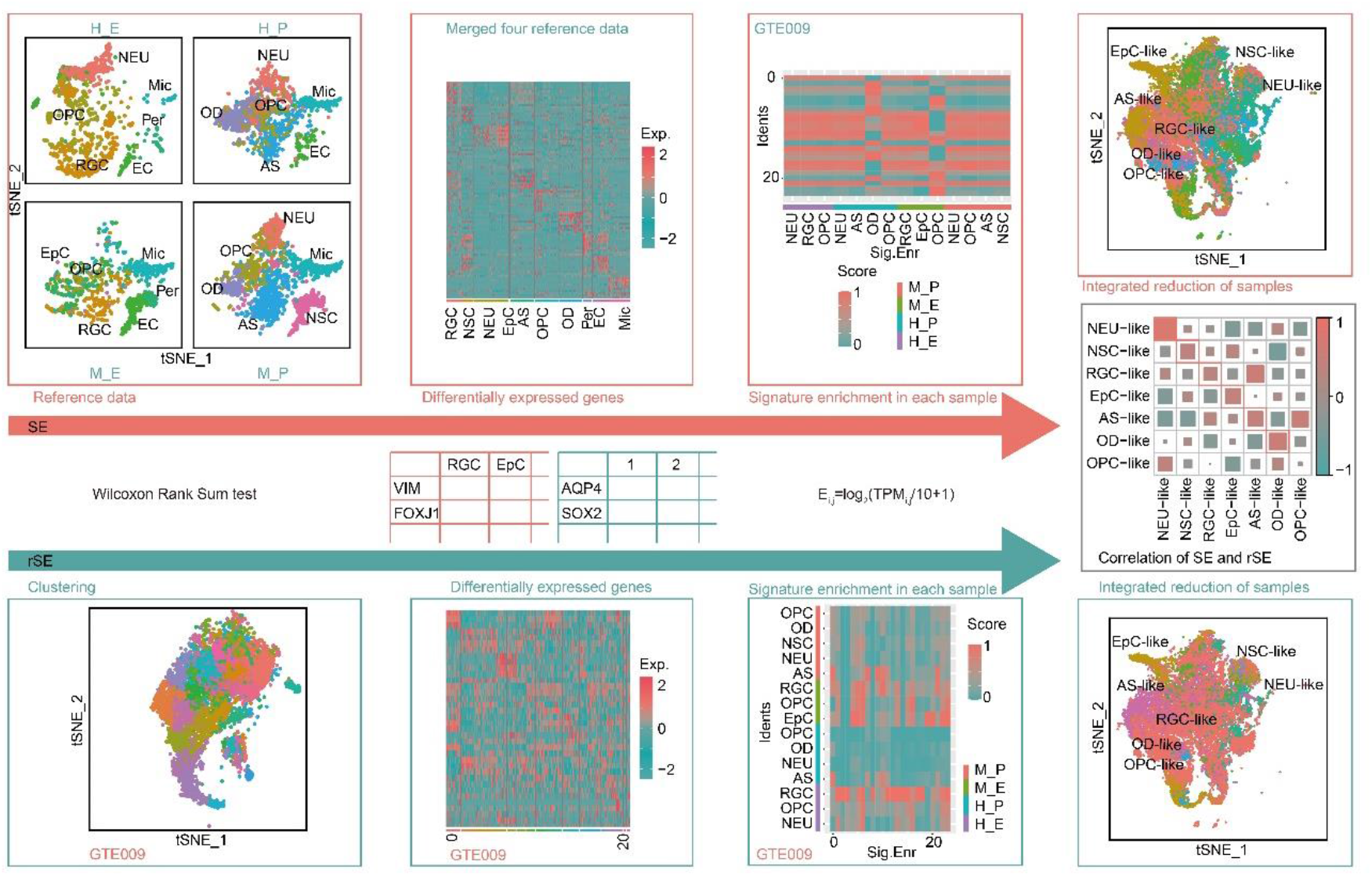
Workflow of Cell Type Classification. Top: Schematic for cell type classification by signature enrichment (SE). The DEGs of different cell populations are obtained from published transcriptomic datasets of human and rodent embryonic and adult cortex (details see Supplementary Table 2) and used as signatures to distinguish cell types. All unknown cells are then clustered at high resolution to obtain multiple clusters, and the highest signature enrichment score in each cluster is designated as the cell type identity for these clusters. Bottom: Schematic for cell type classification by reversed signature enrichment (rSE). In the reciprocal analysis pipeline, unknown cells are first clustered at high resolution to obtain multiple clusters and the DEGs of all clusters are calculated and used as signatures to distinguish cell types. The signatures are then compared with published transcriptomic datasets of human and rodent embryonic and adult cortex (details see Supplementary Table 2), and the highest signature enrichment score is assigned as the name for the unknown cluster. The correlation result of SE- and rSE-determined cell types by ‘cor’ function of stats package in R v3.6.3 confirms high correlation across the two analysis pipelines.

**Supplementary Fig. 4.**
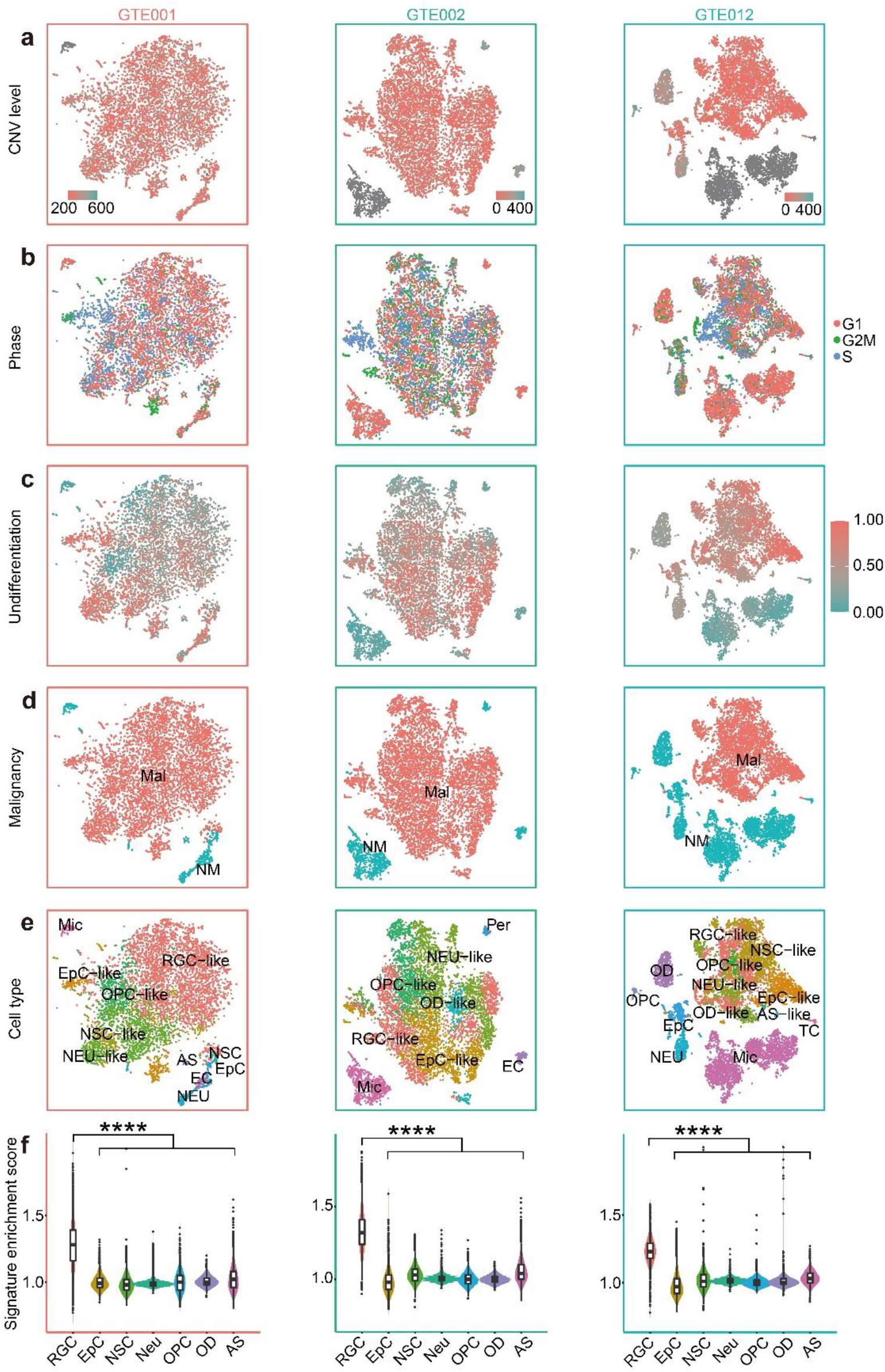
Additional scRNA-Seq Analysis of Human EPN samples. **a**, CNV score calculated by modified inferCNV of samples (GTE001, GTE002, and GTE012) presented on tSNE reduction. **b**, Cell cycle phases in cellular level of samples (GTE001, GTE002, and GTE012) presented on tSNE reduction. **c**, Undifferentiated score calculated by CytoTRACE of samples (GTE001, GTE002, and GTE012) presented on tSNE reduction. **d**, Classified non-malignant cells and malignant tumor cells of samples (GTE001, GTE002, and GTE012) presented on tSNE reduction. **e**, Annotated clusters of samples (GTE001, GTE002, and GTE012) presented on tSNE reduction. **f**, Enrichment of signatures in malignant tumor cells compared to other cell types in samples (GTE001, GTE002, and GTE012; one-way ANOVA analysis; p value < 0.0001). See also Supplementary Table 1.

**Supplementary Fig. 5.**
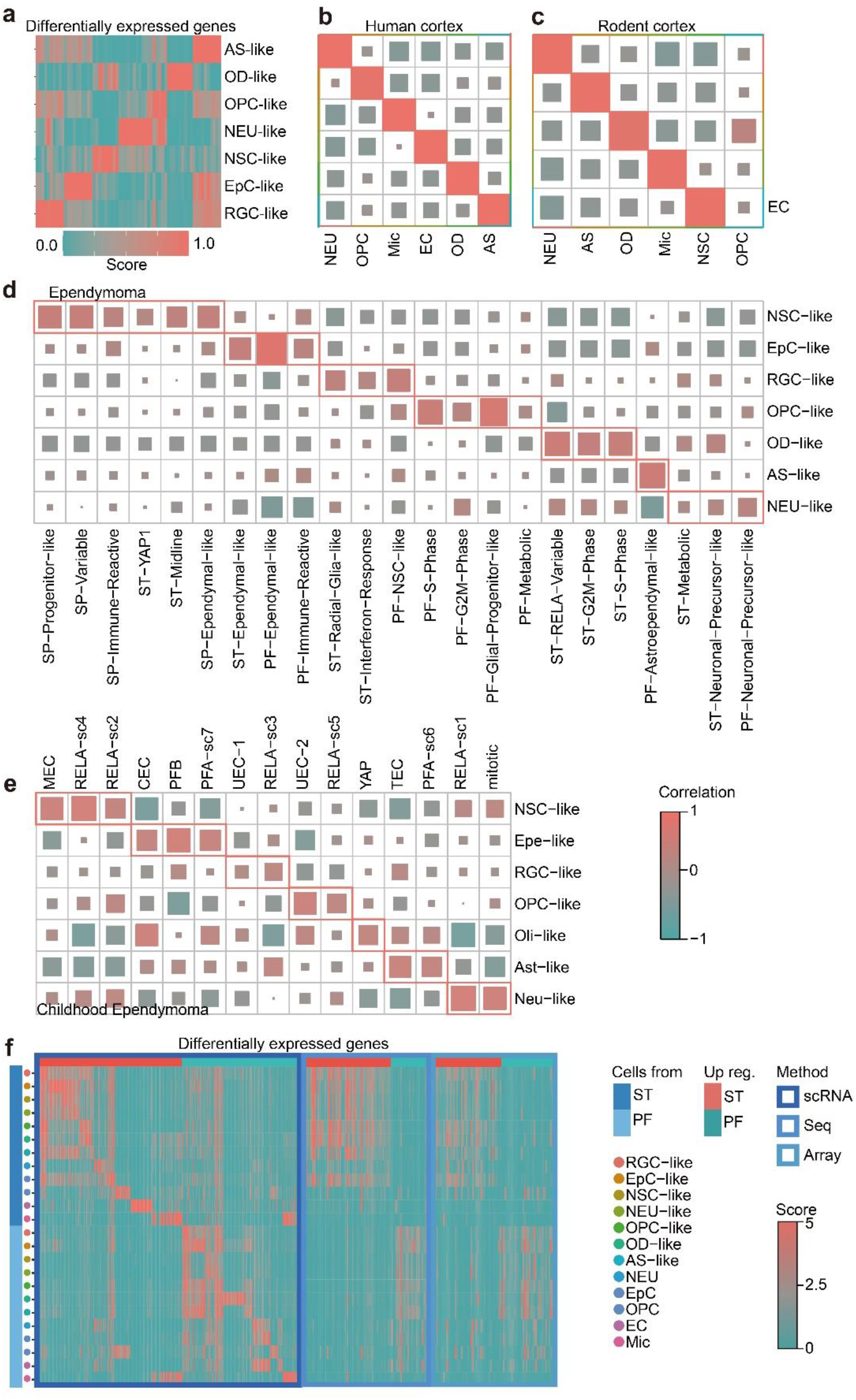
EPN cell type classification validation. **a**, Heatmap of DEGs in annotated clusters of the sample GTE009. **b-e**, Correlation of cell types classified by method of signature enrichment (rows) and that of original cell types determined by original authors (b: human cortex ^46^, c: rodent cortex ^54^, d: ependymoma ^5^, and e: childhood ependymoma ^6^). **f**, Heatmap of DEGs calculated by cell types and pathogenic sites from scRNA-seq data in this study and bulk-DEGs. Aforementioned bulk-DEGs were calculated by pathogenic sites from online bulk-seq data (Gene Expression Omnibus ^55,56^; Seq: GSE89448 ^57^; Array: GSE64415 ^58–60^; aligned to human reference genome GRCh38(hg38)) through DESeq2 ^61^.

**Supplementary Fig. 6.**
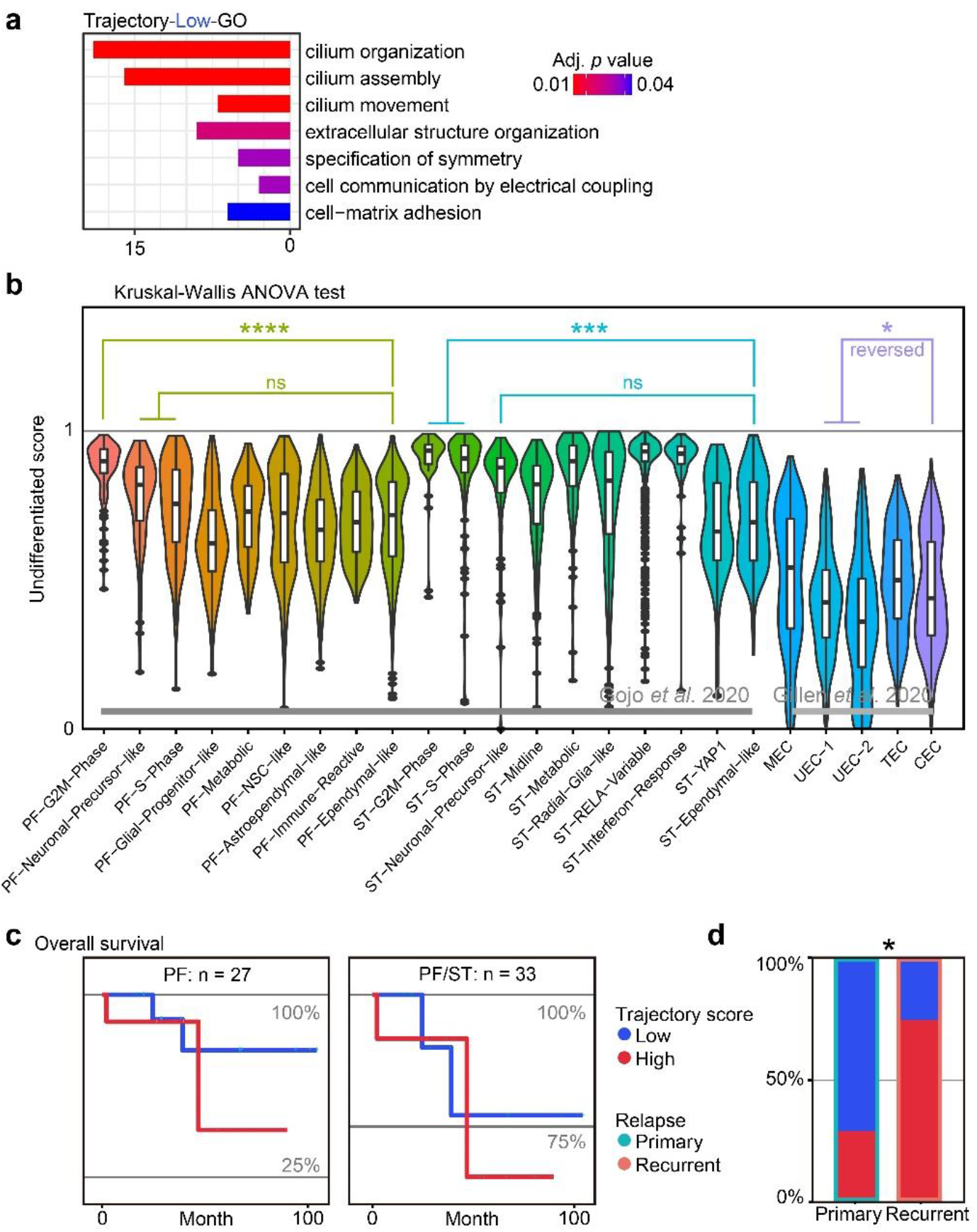
Additional trajectory score analysis of EPN. **a**, Gene ontology analysis of upregulated genes in patients with low trajectory score compared to patients with high trajectory score in published ependymoma scRNA-seq data ^5,6^. **b**, Undifferentiated score analysis on published scRNA-seq datasets of ependymoma ^5,6^ (Kruskal-Wallis test). **c**, Overall survival analysis on the trajectory score. A *p* value of 0.39 and 0.72 were obtained from the difference between two groups of trajectory score of patients (separated by the mean of trajectory score), with a trend of worse survival in patients with high trajectory score. The high and low group were separated by the mean of trajectory score on published scRNA-seq data ^5,6^. **d**, Histogram showing percentage of cells with high and low trajectory score and outlined by subclone annotation in samples from primary and recurrent patients. Permutation test shown significant compositional difference between primary and recurrent samples (*p* value = 0.01241; asymptotic two-sample Fisher-Pitman permutation test). * p < 0.05; ** P < 0.01; *** p < 0.001; **** p < 0.0001

**Supplementary Fig. 7.**
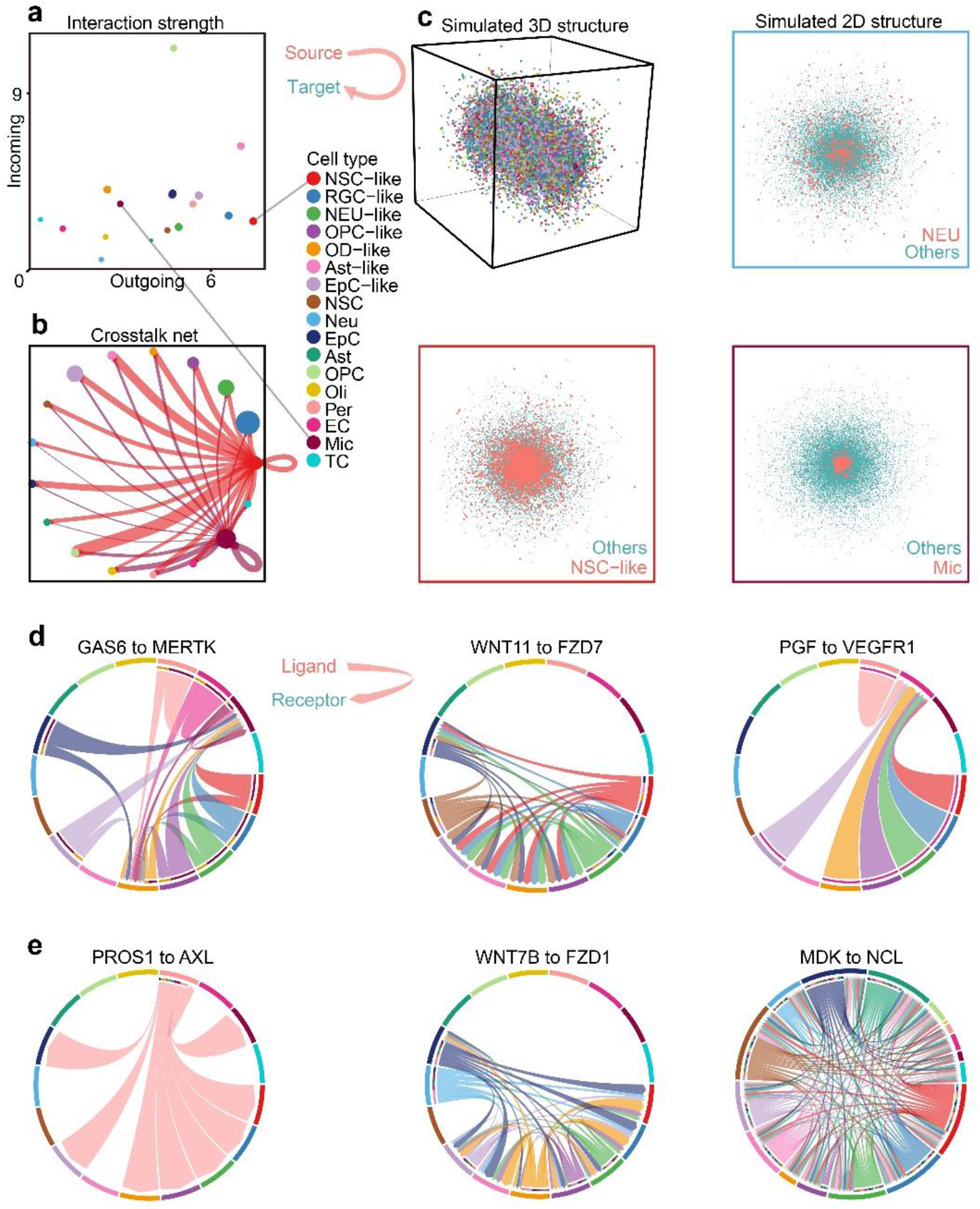
Additional crosstalk analysis of EPN. **a**, Interaction strength between cell types profiled in EPN samples inferred by CellChat. **b**, Crosstalk net analyzed by CellChat. Individual lines represent the crosstalk from source to target cells, highlighting interactions from NSC-like cells and Mic to other cell types. **c**, Simulated 3D spatial structure of all cells and 2D angle of the simulated spatial structure by CSOMAP of Mic, NSC-like cells, and NEU cells respectively, colored by pink (cell type of interest) and blue (other cell types). **d-e**, Circle plots of ligands and receptors with higher expression in recurrent samples than that in primary samples. Lines represent the crosstalk between specific ligands and colors represent the cell type origin for each interaction.

**Supplementary Fig. 8.**
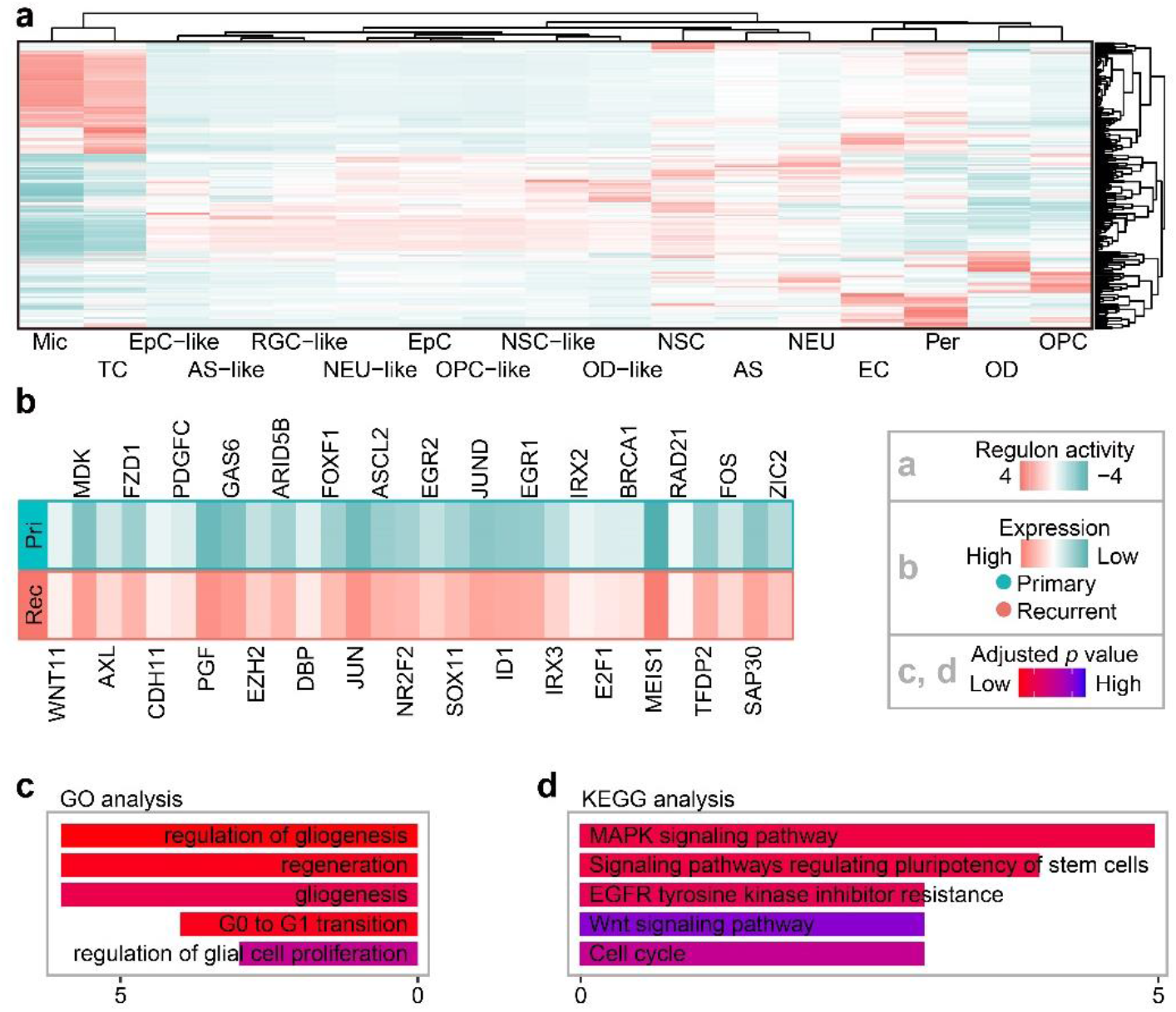
Additional regulon analysis of EPN. **a**, Gene regulatory networks were inferred by SCENIC and were clustered by cell types (bottom) and regulons (right). **b**, List of significantly upregulated genes in recurrent samples from crosstalk and gene regulatory network analysis of NSC-like cells which share the same enriched terms in GO/KEGG analysis. **c-d**, Visualization of genes using GO and KEGG enrichment analysis.

**Table S1. Clinical data from EPN patients.** Clinical data of four sequenced EPN samples from patients in this study.

**Table S2. Signatures for cell type classification.** For recognition of cell type in SE algorithms, Seurat was used to calculate the DEGs (latter utilized as signatures) of each cell type from each reference data 12-16,62 which included RGC, AS, EC, EpC, NEU, NSC, OD, OPC, Mic, and T cells, from human and rodent embryonic and postnatal cortex scRNA-seq data.

**Table S3. Differentially expressed genes between subclones of the sample GTE009.** Differentially expressed genes calculated from subclone 1 compared to the subclone 2 of the sample GTE009.

**Table S4. Analysis on cilium-related genes and CNV score.** DEGs calculated from subclone 1 compared to the subclone 2 of the sample GTE009 marked by GO terms, annotated with results of statistical analysis from CNV score.

**Table S5. Crosstalk analysis output.** List of ligand-receptor interactions annotated by cell type source and target.

## Notes

### Competing Interest Statement

The authors have declared no competing interest.

