## Supplementary figures and images for "Single-cell RNA Sequencing of Pediatric Ependymoma Unravels Subclonal Heterogeneity Associated with Patient Survival"

### Sup. Fig. 1

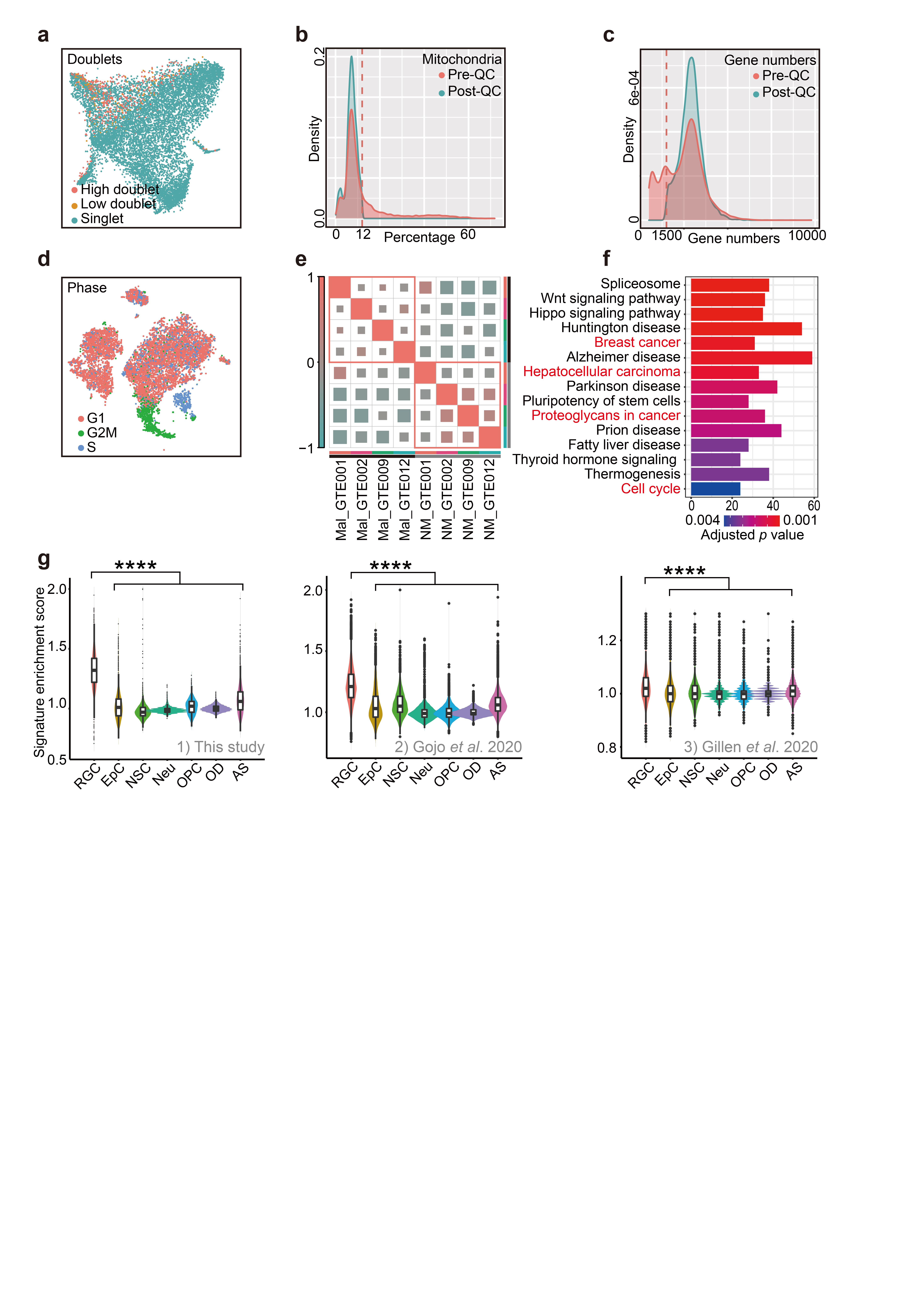

### Sup. Fig. 2

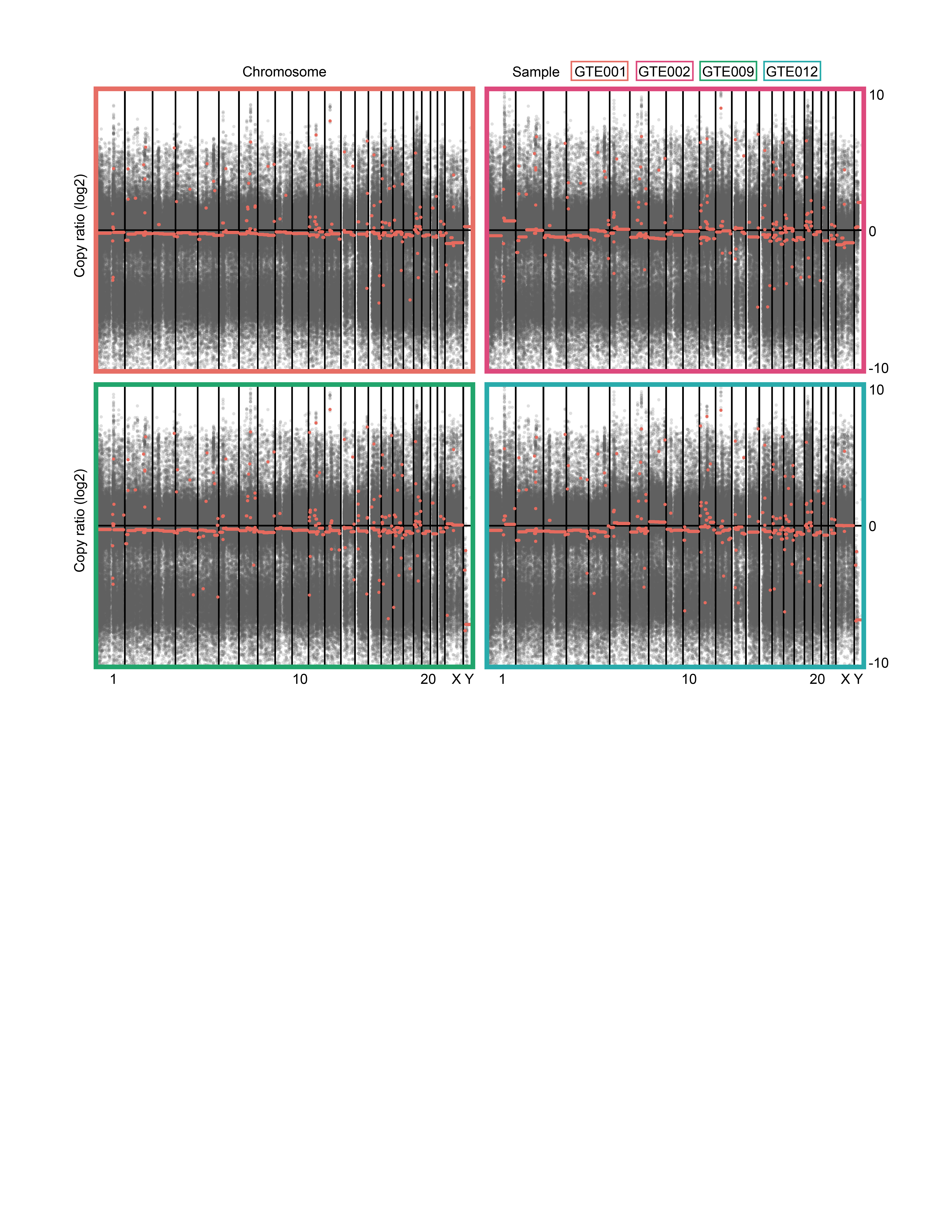

### Sup. Fig. 3

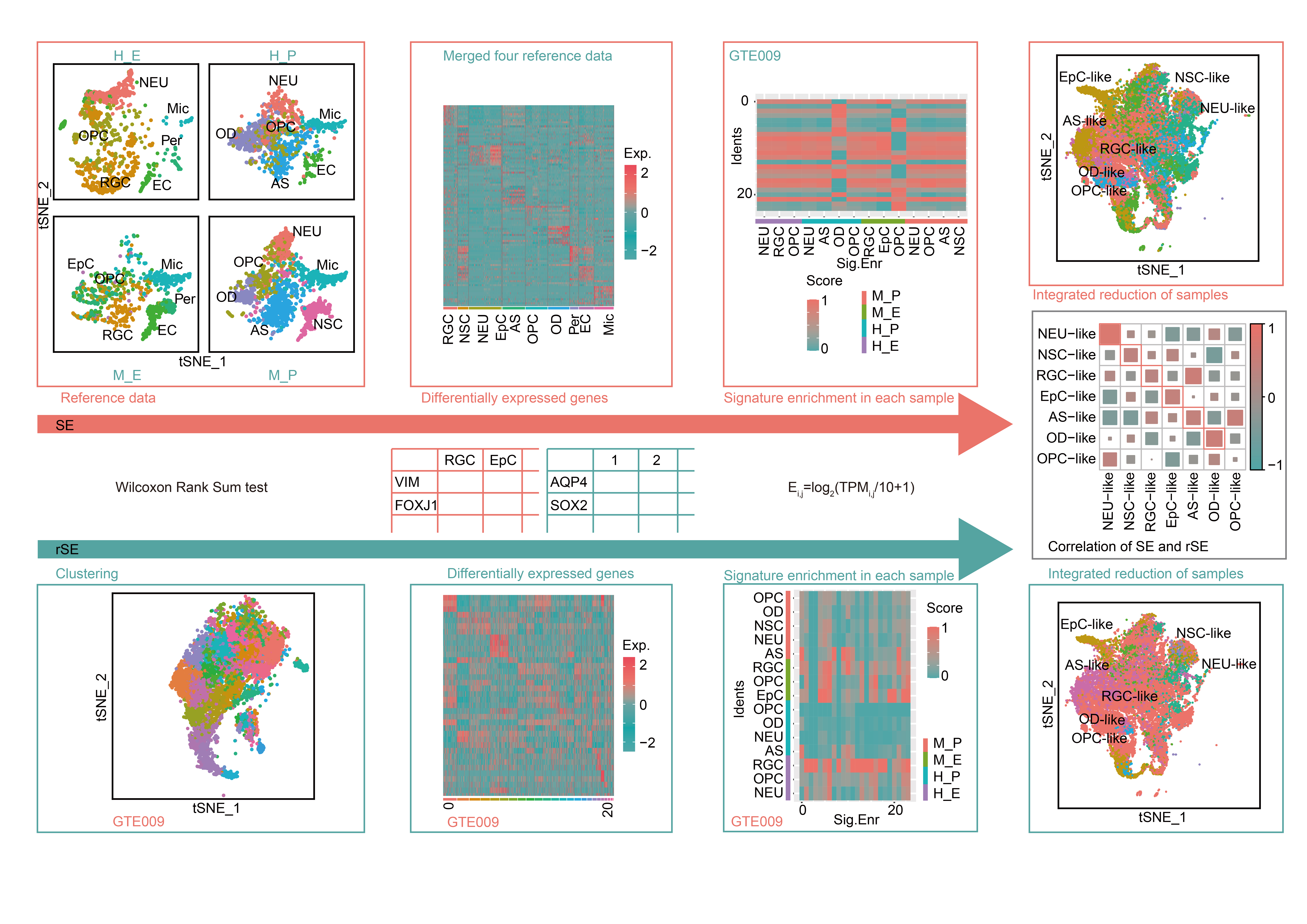

### Sup. Fig. 4

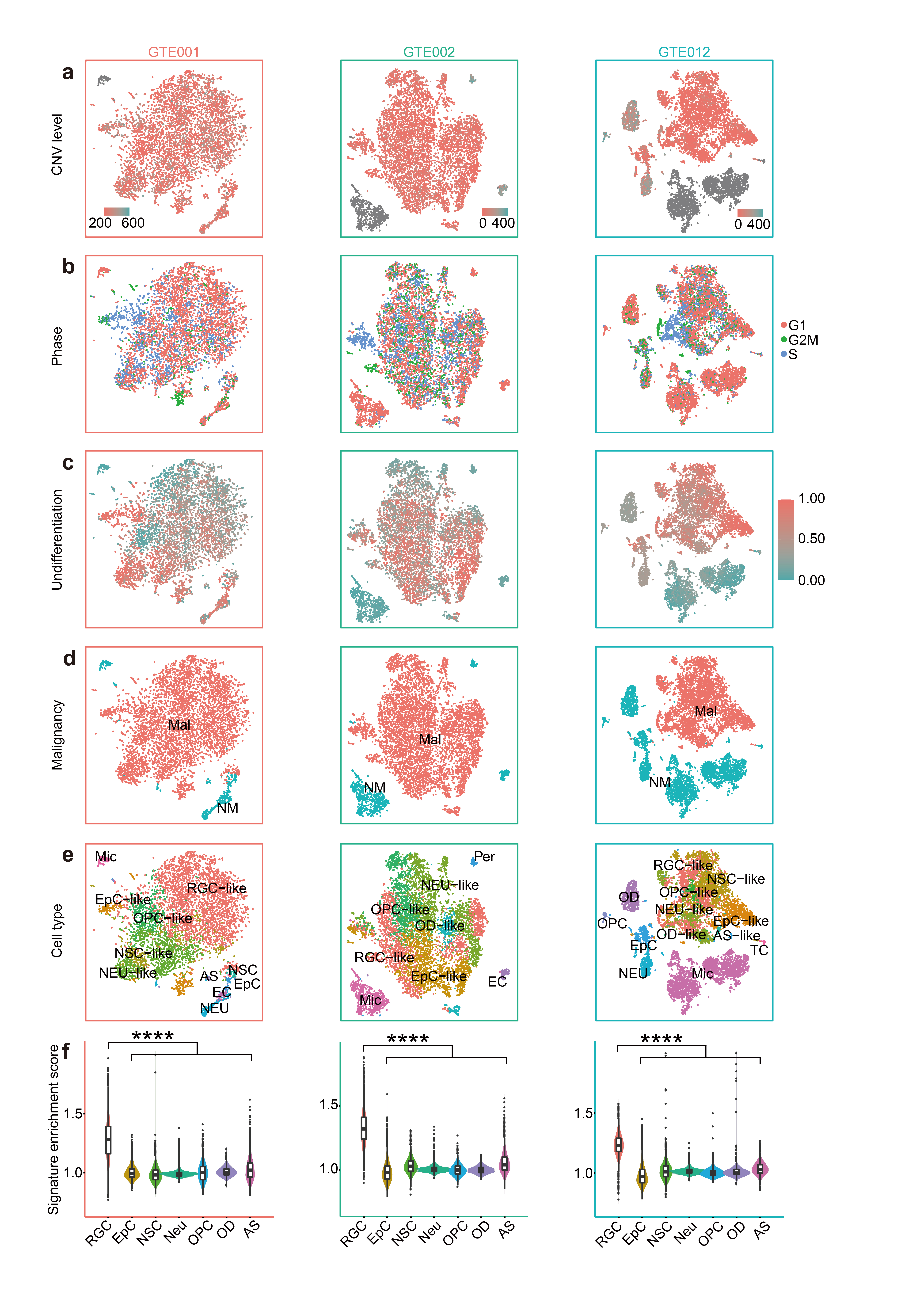

### Sup. Fig. 5

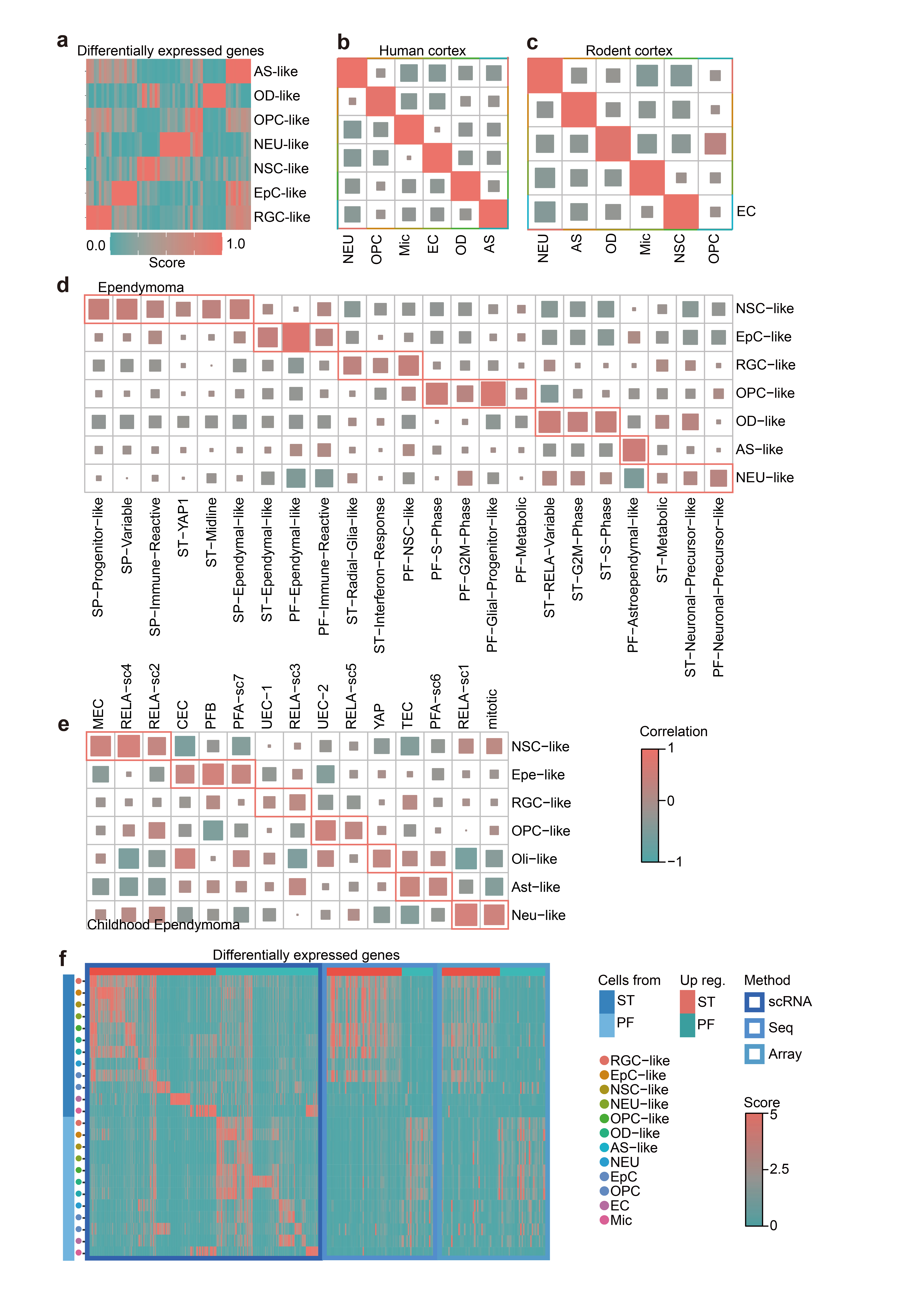

### Sup. Fig. 6

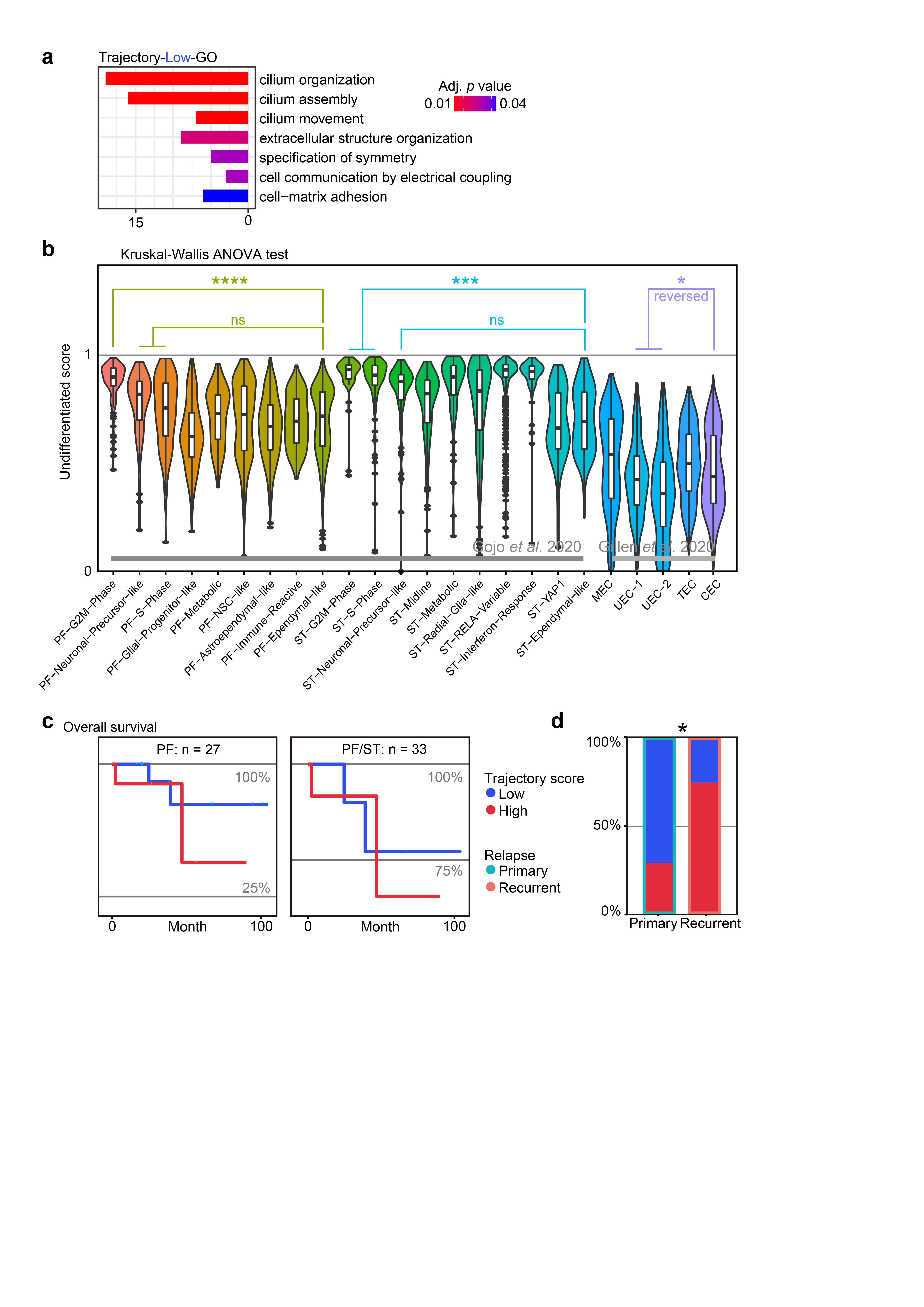

### Sup. Fig. 7

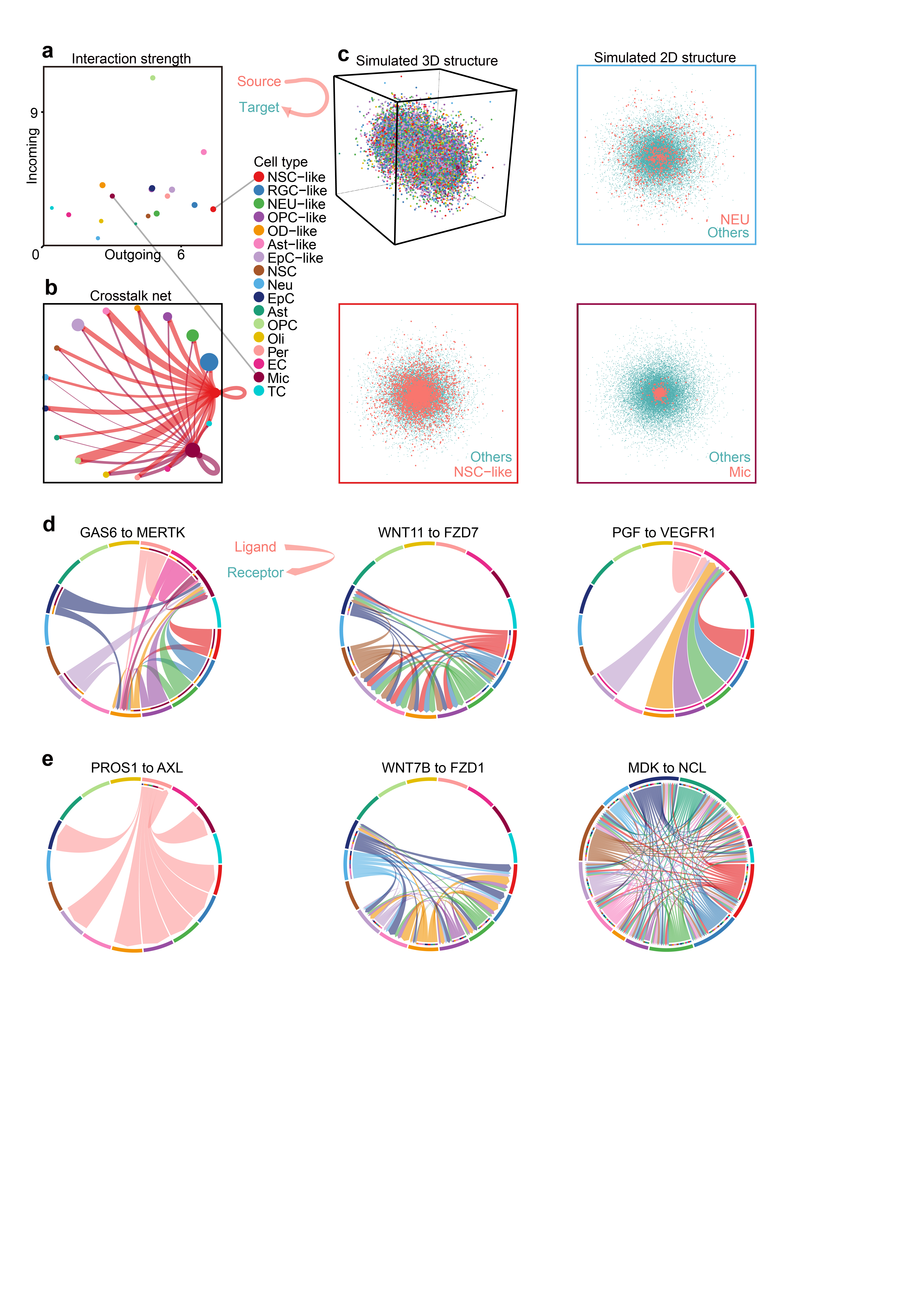

### Sup. Fig. 8

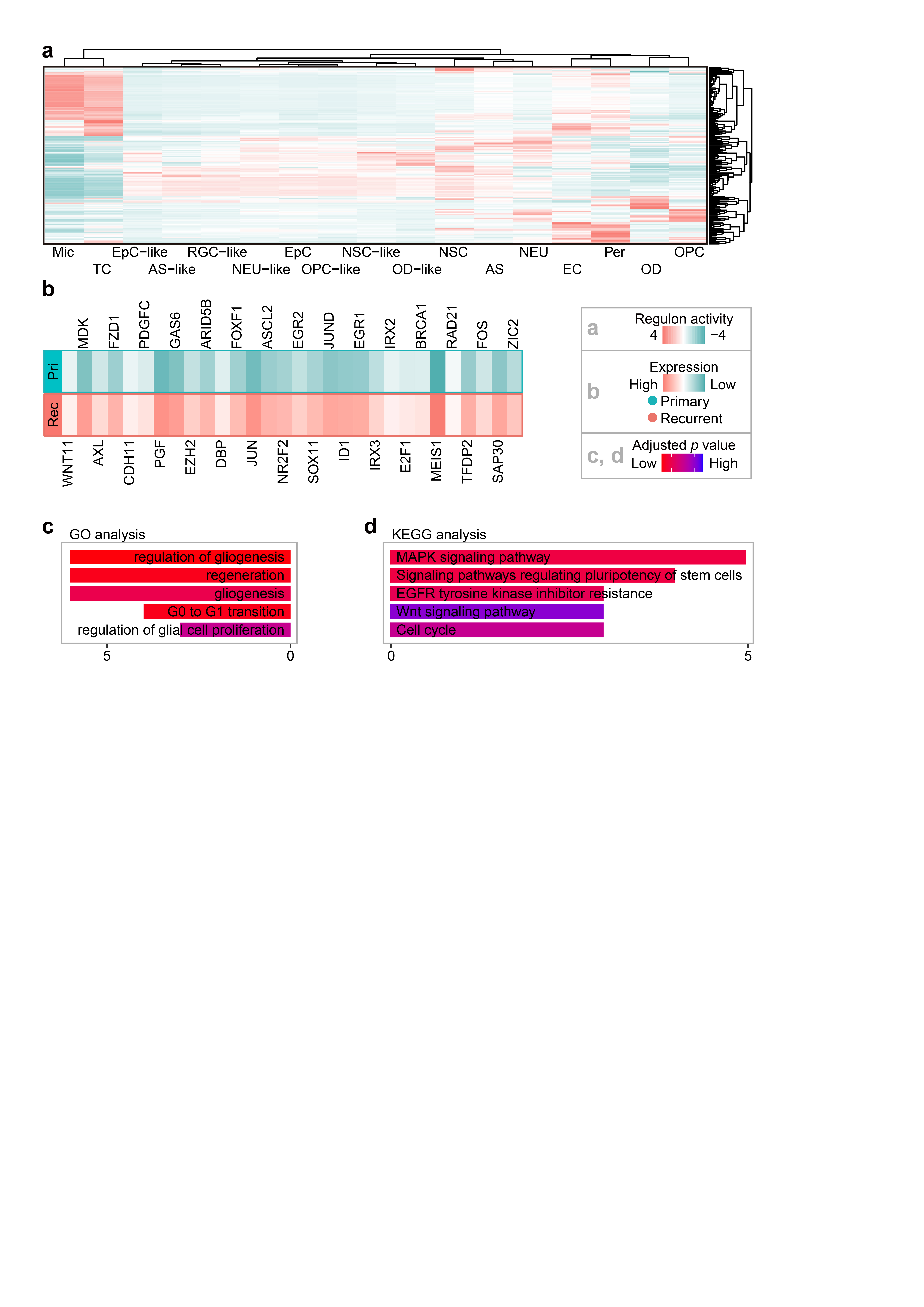
